# The expression profile and tumorigenic mechanisms of CD97 (ADGRE5) in glioblastoma render it a targetable vulnerability

**DOI:** 10.1101/2023.08.31.555733

**Authors:** Niklas Ravn-Boess, Nainita Roy, Takamitsu Hattori, Devin Bready, Hayley Donaldson, Christopher Lawson, Cathryn Lapierre, Aryeh Korman, Tori Rodrick, Enze Liu, Joshua D. Frenster, Gabriele Stephan, Jordan Wilcox, Alexis D. Corrado, Julia Cai, Rebecca Ronnen, Shuai Wang, Sara Haddock, Jonathan Sabio Ortiz, Orin Mishkit, Alireza Khodadadi-Jamayran, Aris Tsirigos, David Fenyö, David Zagzag, Fabiana Perna, Drew R. Jones, Richard Possemato, Akiko Koide, Shohei Koide, Christopher Y. Park, Dimitris G. Placantonakis

**Affiliations:** Department of Neurosurgery, NYU Grossman School of Medicine, New York, NY 10016, USA; Department of Pathology, NYU Grossman School of Medicine, New York, NY 10016, USA; Laura and Isaac Perlmutter Cancer Center, NYU Grossman School of Medicine, New York, NY 10016, USA; Department of Biochemistry and Molecular Pharmacology, NYU Grossman School of Medicine, New York, NY 10016, USA; Metabolomics Laboratory, NYU Grossman School of Medicine, New York, NY 10016, USA; Department of Medicine, Division of Hematology/Oncology, Indiana University, Indianapolis, IN 46202, USA; Preclinical Imaging Laboratory, NYU Grossman School of Medicine, New York, NY 10016, USA; Applied Bioinformatics Laboratories, NYU Grossman School of Medicine, New York, NY 10016, USA; Institute for Systems Genetics, NYU Grossman School of Medicine, New York, NY 10016, USA; Moffitt Cancer Center, Tampa, FL 33612, USA; Department of Medicine, NYU Grossman School of Medicine, New York, NY 10016, USA; Kimmel Center for Stem Cell Biology, NYU Grossman School of Medicine, New York, NY 10016, USA; Brain and Spine Tumor Center, NYU Grossman School of Medicine, New York, NY 10016, USA; Neuroscience Institute, NYU Grossman School of Medicine, New York, NY 10016, USA

**Keywords:** glioblastoma, CD97, adhesion G protein-coupled receptor, Warburg metabolism, antibody-drug conjugate

## Abstract

Glioblastoma (GBM) is the most common and aggressive primary brain malignancy. Adhesion G protein-coupled receptors (aGPCRs) have attracted interest for their functional role in gliomagenesis and their potential as treatment targets. To identify therapeutically targetable opportunities among aGPCR family members in unbiased fashion, we analyzed expression levels of all aGPCRs in GBM and non-neoplastic brain tissue. Using bulk and single cell transcriptomic and proteomic data, we show that CD97 (*ADGRE5*), an aGPCR previously implicated in GBM pathogenesis, is the most promising aGPCR target in GBM, by virtue of its abundance in all GBM tumors and its *de novo* expression profile in GBM compared to normal brain tissue and neural progenitors. CD97 knockdown or knockout significantly reduces the tumor initiation capacity of patient-derived GBM cultures (PDGC) *in vitro* and *in vivo*. Transcriptomic and metabolomic data from PDGCs suggest that CD97 promotes glycolytic metabolism. The oncogenic and metabolic effects of CD97 are mediated by the MAPK pathway. Activation of MAPK signaling depends on phosphorylation of the cytosolic C-terminus of CD97 and recruitment of β-arrestin. Using single-cell RNA-sequencing and biochemical assays, we demonstrate that THY1/CD90 is the most likely CD97 ligand in GBM. Lastly, we show that targeting of PDGCs with an anti-CD97 antibody-drug conjugate *in vitro* selectively kills tumor cells but not human astrocytes or neural stem cells. Our studies identify CD97 as an important regulator of tumor metabolism in GBM, elucidate mechanisms of receptor activation and signaling, and provide strong scientific rationale for developing biologics to target it for therapeutic purposes.

## INTRODUCTION

Glioma is the most common primary brain malignancy. In adults, two main types of glioma exist.^1,2^ The less common type is typically encountered in younger patients and is driven by a neomorphic mutation in the metabolic enzyme isocitrate dehydrogenase (IDH). The more common type, also known as glioblastoma (GBM), is observed in older patients, lacks the IDH mutation (IDH-wildtype), has an aggressive course, and represents the largest unmet need in neuro-oncology. The current treatment regimen for GBM involves neurosurgical resection of the tumor, followed by chemoradiotherapy. Nevertheless, these measures have done little to improve patient outcomes, with the median survival limited to about 15 months.^3-5^ It is clear that in order to improve GBM treatment, we must identify new targetable vulnerabilities of tumor cells and their microenvironment.

In our effort to identify novel mechanisms contributing to GBM tumorigenesis, our group has become interested in the adhesion family of G protein-coupled receptors (aGPCRs), which consists of 32 members.^6^ Adhesion GPCRs are characterized by large extracellular N-termini that contain both receptor-specific domains determining ligand binding and a functionally conserved GPCR autoproteolysis inducing (GAIN) domain, which catalyzes receptor cleavage at the GPCR proteolysis site (GPS).^7-9^ Emerging evidence has implicated several aGPCRs in developmental, physiologic, and oncogenic processes.^6,10-15^ This prompted us to compare the expression of all aGPCR members in cell types within healthy, non-neoplastic brain tissue; neural stem cells (NSCs), the putative cell-of-origin in GBM^16-18^; and patient-derived GBM cultures (PDGCs). Our transcriptomic and proteomic expression analysis identified CD97 (*ADGRE5*) as the aGPCR with the largest differential expression profile: high expression in PDGCs derived from all transcriptional subtypes of GBM in The Cancer Genome Atlas (TCGA; proneural, classical, and mesenchymal)^2,19^, and absence from normal brain tissue and NSCs.

CD97 is expressed in several lineages of the immune system, where it is critical for the inflammatory response,^20-24^ as well as in multiple liquid and solid malignancies.^13,25-28^ Among these malignancies is GBM, in which CD97 was previously implicated in cellular proliferation, brain invasion, and tumor metabolism.^29-33^ However, its mechanism of action in GBM remains incompletely understood. Furthermore, little research effort has been devoted to its therapeutic targeting. Here, we use PDGCs to demonstrate that CD97 is essential for tumor growth both *in vitro* and *in vivo*. Using transcriptomic, metabolomic, and signaling assays, we find that CD97 helps promote glycolytic metabolism via activation of the mitogen-activated protein kinase (MAPK) signaling pathway. We also identify THY1/CD90 as the most likely physiologically relevant CD97 ligand in GBM and demonstrate that CD97 signaling depends on phosphorylation of its C-terminus and recruitment of β-arrestin. To capitalize on CD97’s therapeutic potential, we show that a CD97 antibody-drug conjugate (ADC) that we have developed in-house specifically kills GBM cells, but not human astrocytes or NSCs *in vitro*. Collectively, these data elucidate novel receptor activation and signaling mechanisms employed by CD97 to promote tumor growth and regulate tumor metabolism in GBM, and highlight the receptor’s potential as a therapeutically targetable vulnerability in GBM.

## RESULTS

### CD97 is *de novo* expressed in GBM relative to normal brain

To compare the expression patterns of aGPCRs in GBM relative to normal, non-neoplastic brain tissue, we undertook a combined transcriptomic and proteomic approach. First, we analyzed the mRNA expression levels of all 32 aGPCRs in normal brain tissue using the Allen Brain Map database, and in PDGCs collected in-house (**Figure 1A**). The transcriptional data suggested upregulated expression of several aGPCRs in GBM compared to normal brain cell types. Second, we interrogated proteomic data from normal brain^34^ and GBM (CPTAC - Clinical Proteomic Tumor Analysis Consortium)^35^, and confirmed *de novo* expression at the protein level for some aGPCRs (**Figure 1B**). Third, we compared the differential gene expression of aGPCRs between human NSCs, the putative cell-of-origin in GBM,^18^ and PDGCs using bulk RNA-sequencing (RNA-seq) data (**Figure 1C,D**). Collectively, we found that *ADGRE5* (CD97) fulfilled criteria for the single most differentially expressed aGPCR in GBM compared to healthy brain: it was one of the most upregulated aGPCRs in PDGCs and was detected in all 99 GBM specimens from the CPTAC database^35^; it was absent from non-neoplastic brain tissue at the transcriptomic and proteomic level; and the statistical significance of its transcriptional upregulation in PDGCs compared to NSCs was the highest genome-wide. To confirm these findings, we also performed immunofluorescent staining of normal temporal lobe tissue sections and several (n=10) GBM tissue sections and confirmed that CD97 was uniquely expressed in GBM and not in normal brain (**Figure 1E**). Confocal microscopy confirmed plasma membrane localization of the receptor in tumor cells within GBM biospecimens (**Figure 1F**). Using flow cytometry, we found robust surface expression of CD97 in all tested PDGCs (representing all three TCGA-defined transcriptional subtypes of GBM: proneural, classical, and mesenchymal^2,19^), and low expression in non-neoplastic human astrocytes (NHAs) and NSCs (**Figure 1G; Supp. Fig. 1A**). These data suggest that CD97 is *de novo* expressed in GBM relative to non-neoplastic brain tissue and that it is ubiquitously expressed in all GBM tumors.

**Figure 1:**
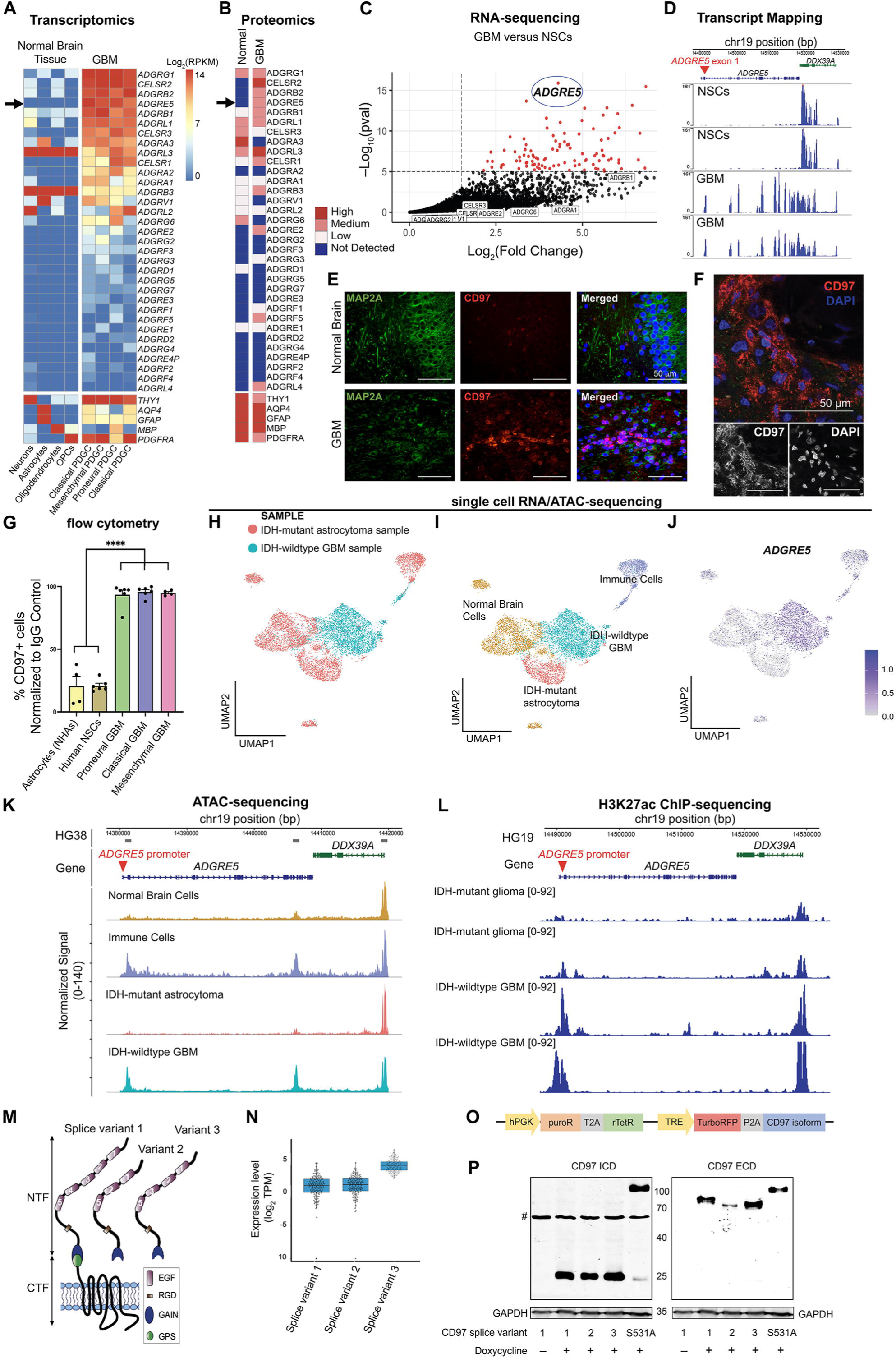
CD97 (*ADGRE5*) is *de novo* expressed in GBM. (**A**) Heatmap showing log_2_ expression levels of aGPCRs and known neural markers in normal brain tissue and GBM [log_2_(RPKM)]. *ADGRE5* is indicated by an arrow. (**B**) Heatmap showing protein expression levels of aGPCRs and known neural markers in normal brain tissue and in GBM. ADGRE5 is indicated by an arrow. “Not detected” indicates expression below the level of mass spectrometry sensitivity. Cut-off parameters for “Low”, “Medium”, and “High” expression are as previously described.^34^ (**C**) Volcano plot showing differentially expressed genes that are upregulated in a PDGC compared to human NSCs. All aGPCRs are labeled. Only *ADGRE5* (circled) was significantly upregulated (p_adj_<0.00001, log_2_(fold change)>1.5). (**D**) Transcripts from bulk RNA-seq data collected from two PDGC replicates and two NSC replicates were aligned along the *ADGRE5* locus. The red arrow signifies the first *ADGRE5* exon. (**E**) Images of immunofluorescent and Hoechst 33342-stained paraffin-embedded specimen of human temporal lobe (normal) and GBM. MAP2A is a neuronal marker. An antibody against the CD97 ECD was used for CD97 staining. (**F**) Confocal microscopy images of a GBM specimen stained with an antibody against the CD97 ECD (red) and DAPI (blue). A merged color image is accompanied by black and white single-channel images. (**G**) CD97 surface staining via flow cytometry using an allophycocyanin (APC)-conjugated antibody in proneural (n=6), classical (n=6), and mesenchymal (n=4) PDGCs, NSCs (n=6), and NHAs (n=4) (unpaired t-test; ****, p<0.0001). (**H**) Single-cell RNA/ATAC-seq data from IDH-mutant astrocytoma and IDH-wildtype GBM surgical specimens. Single cells are color-coded based on their tumor of origin. (**I**) Cluster identities were characterized and named based on their corresponding gene expression profiles. Single cells are colored based on broad cluster identities. (**J**) *ADGRE5* expression in cell clusters based on integrated data. (**K**) ATAC peaks associated with the *ADGRE5* locus were extracted from single-cell ATAC-seq and integrated based on the clusters in **Fig. 1I**. The red arrow signifies the *ADGRE5* promoter. Also shown are peaks corresponding to the *DDX39A* promoter, which is expressed in all cell clusters. (**L**) H3K27ac ChIP-seq from the GEO shows peaks associated with the *ADGRE5* locus for IDH-wildtype GBM and IDH-mutant anaplastic astrocytoma. The red arrow signifies the *ADGRE5* promoter. (**M**) Schematic representation of the structure of three CD97 isoforms. EGF, epidermal growth factor-like repeat; RGD, arginylglycylaspartic acid domain; GAIN, GPCR autoproteolysis inducing domain; GPS, GPCR proteolysis site. Note that for isoforms 2 and 3, only the NTF is shown. (**N**) Bulk RNA-seq data from the GBM dataset of TCGA were used to quantify CD97 isoform expression. Blue boxes are centered around the median and encompass the two middle quartiles. (**O**) Schematic representation of the tet-inducible vector used for CD97 isoform overexpression. (**P**) Immunoblots for the CD97 ICD (equivalent to the CTF) and the ECD (equivalent to NTF). CD97 S531A is an uncleavable point mutant of isoform 1. Non-specific bands are indicated by a # sign. The same membrane was stained for CD97 ICD and then stripped for CD97 ECD staining. Error bars throughout this figure indicate SEM.

We then tested whether *ADGRE5* expression differs between IDH-wildtype GBM and low-grade IDH-mutant gliomas using single-cell RNA/ATAC (Assay for Transposase-Accessible Chromatin)-seq data collected from an IDH-mutant WHO (World Health Organization) grade II astrocytoma and an IDH-wildtype GBM (**Figure 1H**). After identifying cell clusters based on single-cell RNA-seq gene expression (**Figure 1I; Supp. Fig. 1B**) via Seurat, we found that *ADGRE5* expression was highest in IDH-wildtype GBM tumor cells and immune cells, but substantially less abundant in IDH-mutant low-grade astrocytoma cells (**Figure 1J**). Analysis of single-cell ATAC-seq data using Signac demonstrated open, accessible chromatin at the *ADGRE5* locus only in GBM and immune cell clusters, but not IDH-mutant astrocytoma cells and normal brain lineages (**Figure 1K**). Similarly, chromatin immunoprecipitation (ChIP)-seq for H3K27 acetylation, an activating chromatin modification, obtained from publicly available Gene Expression Omnibus (GEO) datasets^36^, showed increased activation around the *ADGRE5* promoter in IDH-wildtype GBM but not in IDH-mutant gliomas (**Figure 1L**). Together, these findings suggest that CD97 expression is abundant in IDH-wildtype GBM but only limited in low-grade IDH-mutant astrocytoma.

Alternative splicing of *ADGRE5* mRNA generates three distinct isoforms that differ in the number of epidermal growth factor (EGF)-like repeats in the N-terminal extracellular domain (ECD) of the receptor (**Figure 1M**).^25^ Analysis of TCGA RNA-seq data indicated that isoform 3, with four EGF repeats in the N-terminus, is the predominant variant in GBM (**Figure 1N**). CD97 transcript was found in all transcriptional subtypes of GBM in TCGA (**Supp. Fig. 1C**). The extent of autoproteolytic cleavage for each aGPCR sometimes varies depending on the cellular context and cell type.^37^ To test whether CD97 undergoes cleavage at the GPS site in GBM, we exogenously expressed all three isoforms of CD97 and a non-cleavable point mutant of CD97 (CD97 S531A)^38^ in PDGCs (**Figure 1O**). Immunoblotting against the ECD and the intracellular domain (ICD) of CD97 revealed that the vast majority of the receptor undergoes cleavage in PDGCs to generate a membrane-bound C-terminal fragment (CTF) and an extracellular N-terminal fragment (NTF), regardless of the isoform that is overexpressed (**Figure 1P**). To resolve discrepancies in the expected and observed molecular weights of the NTF (**Supp. Fig. 1D**), we deglycosylated whole cell lysates collected from a PDGC endogenously expressing CD97 or exogenously overexpressing the CD97 isoforms before immunoblot staining. The deglycosylation resulted in band shifts to the expected molecular weights (**Supp. Fig. 1E**), suggesting that the CD97 NTF is glycosylated. Interestingly, the band corresponding to endogenously-expressed deglycosylated CD97 NTF appears to match the molecular weight of the NTF of exogenously-expressed CD97 isoform 3 (**Supp. Fig. 1E**). This finding supported the transcriptomic TCGA data indicating that the predominant species of CD97 in GBM is isoform 3 (with four extracellular EGF repeats) and suggested that CD97 undergoes autoproteolytic cleavage in GBM.

### CD97 loss compromises GBM growth

To further investigate the role of CD97 in GBM, we established short hairpin RNAs (shRNAs) and guide RNAs (gRNAs) targeting all isoforms of CD97 (**Figure 2A; Supp. Fig. 2A**). Flow cytometric surface staining confirmed loss of CD97 upon shRNA-mediated knockdown (**Figure 2B**) and CRISPR-Cas9-mediated knockout (**Supp. Fig. 2B**). Since CD97 is implicated in GBM cell migration^29,32,33^, we performed a transwell migration assay which confirmed significant reduction in PDGC migration after lentiviral infection with the CD97 shRNA compared to a scrambled (SCR) shRNA control (**Supp. Fig. 2C**). Importantly, knockdown of CD97 produced a substantial growth defect in our PDGCs *in vitro*. This was quantified using tumorsphere formation assays, which revealed a significant loss of tumorsphere formation after CD97 knockdown (**Figure 2C,D**). Overexpression of a doxycycline (dox)-inducible shRNA-resistant form of *CD97* cDNA alone did not increase tumorsphere formation in PDGCs (**Supp. Fig. 2D**), but rescued the impairment in tumorsphere formation following CD97 knockdown (**Figure 2C,D**), suggesting specificity of the phenotype observed with CD97 shRNA. The effects of CD97 knockdown on PDGCs were also observed in extreme limiting dilution assays (**Figure 2E,F**), suggesting a diminished clonogenic frequency and impaired GBM stem cell self-renewal in the absence of CD97. We also observed significant effects of CD97 on cellular proliferation using cell cycle analysis (**Supp. Fig. 2E**) and Ki67 immunofluorescent staining (**Supp. Fig. 2F**), as well as on cellular viability using flow cytometry for Annexin-V and DAPI (4’,6-diamidino-2-phenylindole) staining (**Supp. Fig. 2G**). In contrast, CD97 knockdown in NHAs did not compromise their proliferation or their viability (**Supp. Fig. 2E**,G). CRISPR-Cas9 knockout of CD97 phenocopied the shRNA knockdown, and significantly hindered PDGC growth and viability relative to a gRNA directed at the safe harbor human *ROSA26* locus in a cell competition assay (**Figure 2G**). To determine whether CD97 knockdown also impaired tumor formation *in vivo*, intracranial xenograft assays were performed in immunodeficient NSG mice. Implantation with PDGCs following CD97 knockdown resulted in significantly less tumor growth (**Figure 2H-J**) and significantly prolonged mouse survival (**Figure 2K**). These findings demonstrate that CD97 loss compromises PDGC proliferation and viability and reduces tumor initiation *in vitro* and *in vivo*.

**Figure 2:**
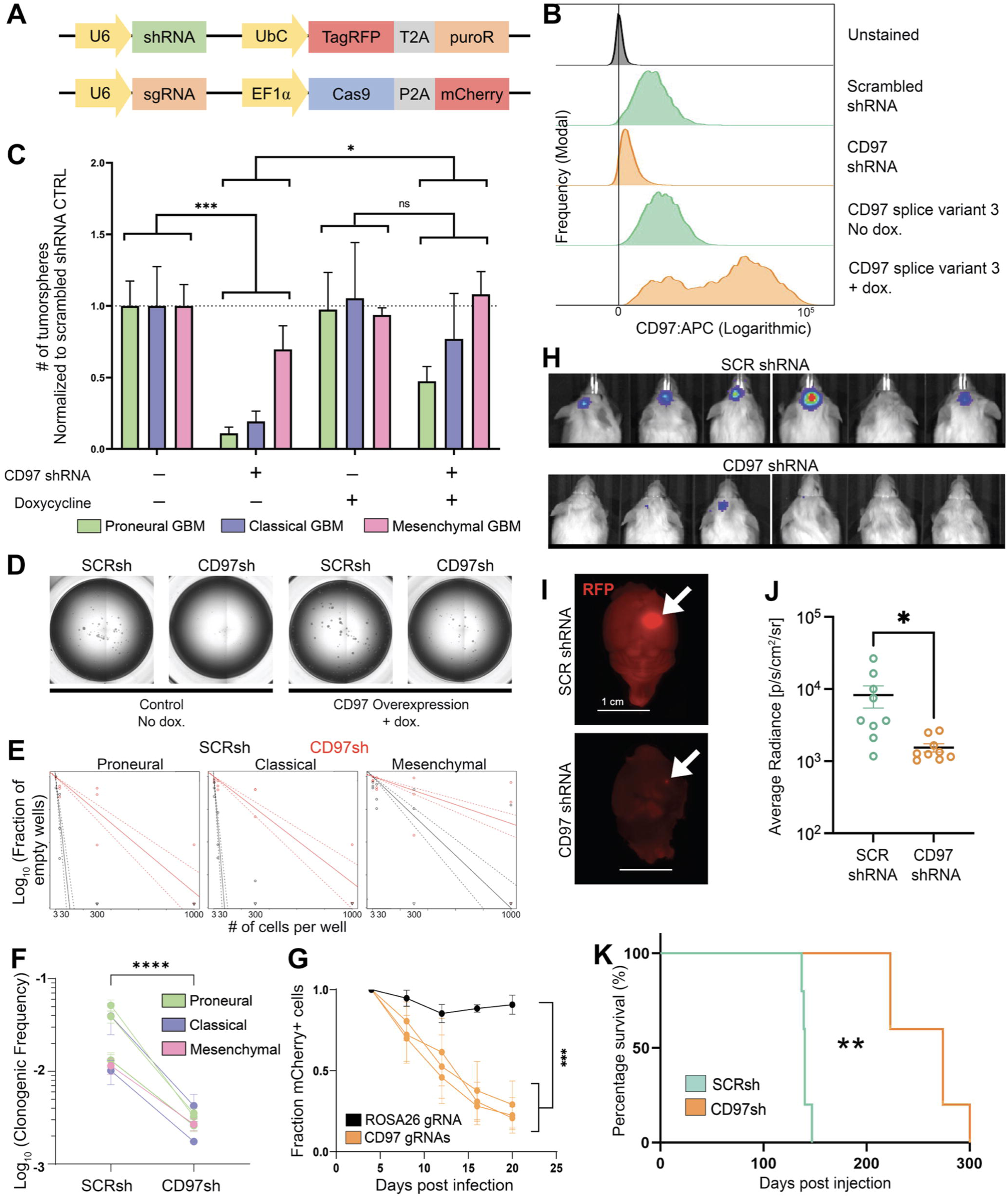
CD97 is essential for tumor initiation *in vitro* and *in vivo*. (**A**) Schematic representations of the shRNA and CRISPR constructs used to knock down and knock out CD97, respectively. (**B**) Histograms of CD97 surface staining following knockdown or overexpression of isoform 3 of CD97 in a PDGC. (**C**) Bar graph visualizing tumorsphere formation following knockdown of CD97 and followed by rescue with a dox-inducible and a shRNA-resistant form of CD97 isoform 3 (n=5 for each PDGC; 2-way ANOVA F_3,36_=9.795, p<0.0001; Tukey’s multiple comparisons test: EV SCRsh vs. EV CD97sh: ***, p<0.001; EV CD97sh vs. OE CD97sh: *, p<0.01; OE SCRsh vs. OE CD97sh: ns, p>0.05.). (**D**) Representative wells from the tumorsphere formation assay quantified in **Fig. 2C** before and after dox-induced rescue. (**E**) ELDA plots for three PDGCs are displayed. (**F**) Summary plot showing calculated clonogenic frequencies for all tested PDGCs after CD97 knockdown based on ELDA assays. Paired samples are connected with a line. (n=3 per PDGC; 2-way ANOVA F_1,28_=75.24, p<0.0001). (**G**) Graph showing compromised viability of PDGCs infected with three separate gRNAs against CD97 (orange) compared to a control gRNA against the human homologue of *ROSA26* (black) in a cell competition assay (n=3 per gRNA; 2-way ANOVA F_4,40_=10.37, p<0.001; Tukey’s multiple comparisons test: ROSA26 vs. CD97: D4, D8, D12: ns, p>0.05; D16, D20: ***, p<0.001). (**H**) Bioluminescent images taken with IVIS of intracranial GBM xenografts 90 days after injection. (**I**) Immunofluorescent images of mouse brains injected with PDGCs lentivirally infected with the indicated shRNA construct, the TagRFP fluorophore, and luciferase. White arrows indicate the resulting tumor at the site of injection. (**J**) Graph depicting the average bioluminescent radiance captured by IVIS for mice harboring PDGC xenografts (n=9 mice per group; unpaired t-test; *, p<0.05). (**K**) Kaplan-Meier survival curve showing significantly increased survival of mice xenografted with a PDGC harboring CD97 shRNA (n=5 mice per group; logrank test; **, p<0.01). Error bars throughout this figure indicate SEM.

### CD97 knockdown impairs glycolytic metabolism

To elucidate mechanisms underlying the actions of CD97 in PDGCs, bulk RNA-seq was performed following knockdown of the receptor in one of our PDGCs. In order to capture early consequences of CD97 knockdown before any effects on cellular viability, we collected cells four days after infection with the lentiviral shRNA vector. Over 3000 genes were differentially expressed between the CD97 knockdown and the SCR shRNA control PDGCs (**Figure 3A**). Among the most downregulated genes was *ADGRE5*, confirming a robust knockdown of the receptor (**Figure 3A**). Gene Ontology (GO) PANTHER pathway analysis to detect gene sets that were depleted upon CD97 knockdown identified glycolytic metabolism as the most impacted biological process (**Figure 3B**). This is noteworthy, because GBM tumor cells exhibit increased reliance on glycolytic metabolism, a phenomenon known as the Warburg effect, compared to normal brain cells. ^39-42^ This led us to hypothesize that CD97 plays a role in promoting glycolysis in GBM.

**Figure 3:**
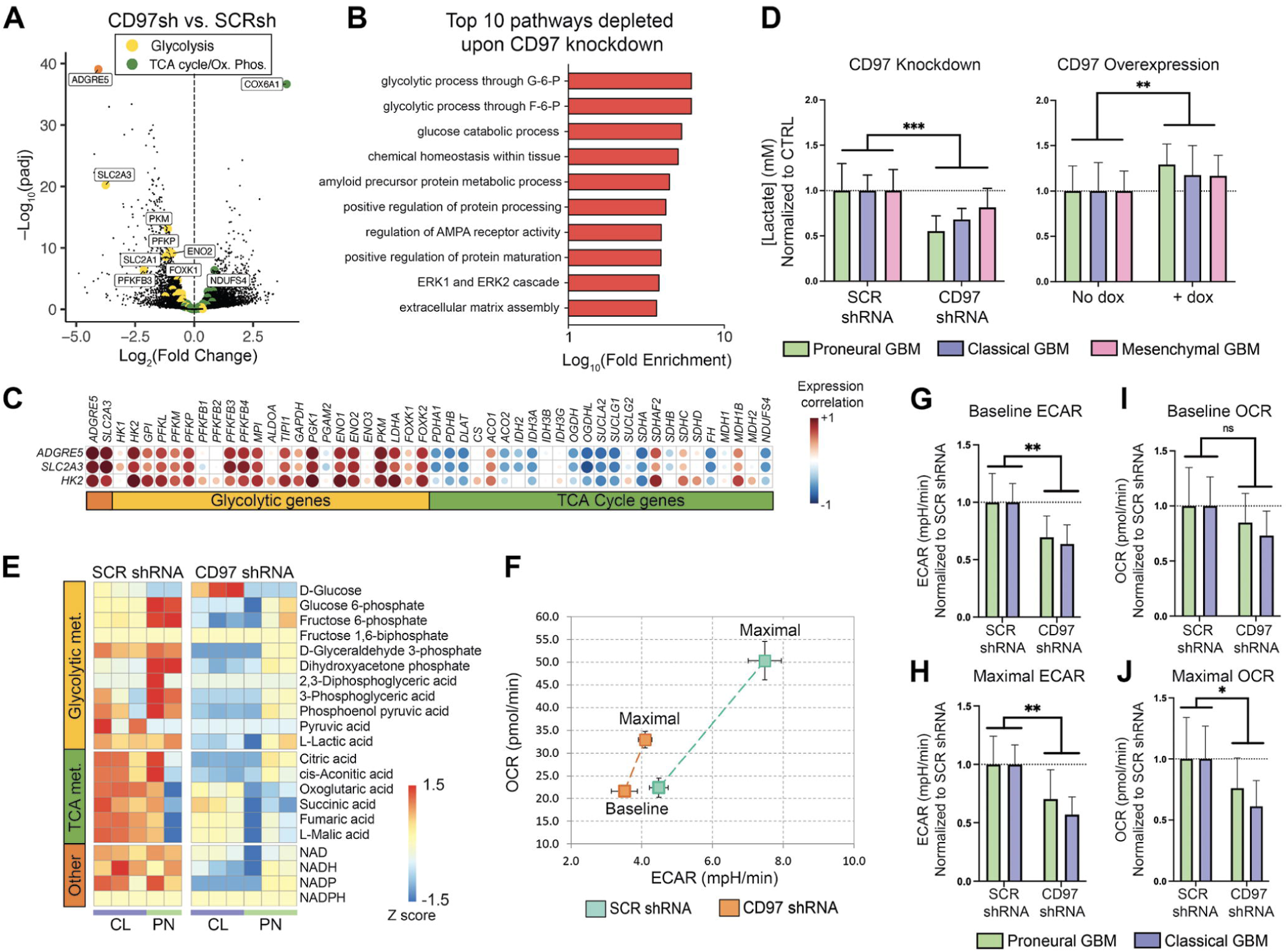
CD97 knockdown impairs glycolytic metabolism in GBM. (**A**) Volcano plot derived from bulk RNA-seq data collected from PDGCs lentivirally infected with the scramble (SCR) or CD97 shRNA (n=3 biological replicates for each). Genes involved in canonical glycolysis and glucose transport are represented as large yellow points. Genes involved in the TCA cycle and OXPHOS (cytochrome oxidase subunits) are represented as large green points. *ADGRE5* (CD97) is represented by an orange point. (**B**) The top ten pathways with the largest fold enrichment among downregulated genes using GO PANTHER pathway enrichment analysis. (**C**) Correlation matrix from bulk RNA-seq data collected from PDGCs following knockdown or overexpression of CD97 shows high correlation between *ADGRE5* and glycolysis-related genes and anti-correlation with TCA cycle-related genes. *HK2* (hexokinase 2) and *SLC2A3* (glucose transporter 3/GLUT3) transcripts are both included because they have been implicated in Warburg metabolism. (**D**) Bar graph showing decreased lactate production after knockdown of CD97 (n=6 per PDGC; 2-way ANOVA F_1,15_=27.24, p<0.001) and increased lactate production after CD97 overexpression in PDGCs (n=3 per PDGC; 2-way ANOVA F_1,6_=26.07, p<0.01). (**E**) Steady-state metabolomic data reveals depletion of glycolytic metabolites after knockdown of CD97 in PDGCs [PN = proneural, n=2-3 (one replicate was removed for technical reasons); CL = classical, n=3]. (**F**) Representative Seahorse Cell Energy Phenotype graph showing OCR and ECAR changes before (baseline) and after (maximal) addition of mitochondrial stressors. (**G, H**) Bar graphs quantifying baseline and maximal ECAR (n=6 per PDGC; Baseline: 2-way ANOVA F_1,10_=14.06; **, p<0.01; Maximal: 2-way ANOVA F_1,10_=17.87, p<0.01). (**I, J**) Bar graphs quantifying baseline and maximal OCR (n=6 per PDGC; Baseline: 2-way ANOVA F_1,10_=2.820, p>0.05; Maximal: 2-way ANOVA F_1,10_=5.474; *, p<0.05). Error bars throughout this figure indicate SEM.

Indeed, when we revisited the differential gene expression data, we found that the majority of genes involved in glycolysis and glucose transport were downregulated upon CD97 knockdown, while the opposite was observed for genes involved in the tricarboxylic acid (TCA) cycle and in oxidative phosphorylation (OXPHOS) (**Figure 3A**). In order to further visualize this, we performed Differential Gene Correlation Analysis (DGCA) of our transcriptomic data from a PDGC following both overexpression or knockdown of CD97 (**Figure 3C**). This analysis also revealed that *ADGRE5* expression levels positively correlated with many genes relevant to glycolysis and negatively correlated with genes relevant to the TCA cycle (**Figure 3C**). We also confirmed a reduction in the glucose transporter GLUT3 (*SLC2A3*) after CD97 knockdown at the protein level via immunoblot staining (**Supp. Fig. 3A**). Of note, several glycolytic enzymes and glucose transporters have been previously implicated in GBM pathogenesis^42-45^ and metabolic reprogramming.^46-48^ In order to ensure that the effects of CD97 knockdown on the transcription of metabolic genes were generalizable, we performed Gene Set Enrichment Analysis (GSEA) on RNA-seq data collected from three other PDGCs following CD97 knockdown. Here, we also observed a depletion of glycolytic processes and an enrichment of OXPHOS-related transcripts following CD97 knockdown (**Supp. Fig. 3B,C**).

Based on the transcriptomic data, we decided to investigate the metabolic consequences of CD97 perturbation in our PDGCs. Since Warburg metabolism is characterized by glucose catabolism biased toward lactate generation rather than pyruvate conversion to acetyl-CoA for utilization in the TCA cycle, we measured lactate levels in the culture medium after shRNA-mediated silencing or overexpression of CD97 in our PDGCs (**Figure 3D**). Indeed, CD97 knockdown resulted in reduced levels of extracellular lactate, while levels increased upon CD97 overexpression. We next performed both steady-state and flux metabolomic analysis in order to gain further insight on the regulation of enzymatic steps of glucose metabolism (**Figure 3E**, **Supp. Fig. 3D**). Steady-state metabolomic data showed a depletion of glycolytic metabolites following CD97 knockdown. Given that glycolytic intermediates feed into both the TCA cycle and the pentose-phosphate pathway (PPP), which is essential for nucleic acid synthesis and redox homeostasis, it is no surprise that these metabolites were also depleted after knockdown (**Figure 3E; Supp. Fig. 3E**).^49^

Metabolic flux data generated using [U-^13^C_6_]-glucose also revealed reduced levels of heavy carbon incorporation into glycolytic metabolites, while the effects on TCA cycle metabolites were less evident (**Supp. Fig. 3D**). Collectively, the transcriptomic and metabolomic data suggest that CD97 knockdown leads to a global reduction in glucose catabolism, which is likely to account for the reduced proliferation and viability of GBM cells. We postulate that the upregulation of transcripts encoding enzymes of the TCA cycle and OXPHOS reflects inadequate compensation aimed to meet energy demands impacted by reduced glycolysis.

To further test the hypothesis that CD97 promotes glycolytic metabolism at a functional level, we measured rates of glycolysis and mitochondrial respiration using a Seahorse XF Cell Energy Phenotype assay. This assay measures the extracellular acidification rate (ECAR), largely accounted for by lactate production, and the oxygen consumption rate (OCR) before and after the addition of mitochondrial stressors (oligomycin and FCCP). PDGCs following CD97 knockdown exhibited a loss of the ability to respond to mitochondrial stressors (**Figure 3F**). Closer investigation revealed a significantly decreased ECAR both at baseline and after the addition of mitochondrial stressors (maximal ECAR) (**Figure 3G,H**). Though the OCR levels were also decreased following CD97 knockdown, these effects were not as striking as those with ECAR and did not reach statistical significance at baseline (**Figure 3I,J**). Together, these data suggest that CD97 promotes glycolytic metabolism in GBM.

### CD97 activates the MAPK/ERK signaling pathway

In an effort to identify signaling pathways activated by CD97 in GBM, we revisited the GO PANTHER pathway analysis of our RNA-seq data shown previously (**Figure 3B**). Here, transcripts involved in the ERK1/ERK2 signaling cascade were depleted upon CD97 knockdown in our PDGCs. To confirm whether the MAPK pathway is activated by CD97 in our PDGCs, we performed immunoblot staining against phosphorylated ERK1/ERK2 following CD97 knockdown or overexpression. CD97 knockdown significantly decreased levels of phosphorylated ERK1/ERK2 (**Figure 4A,B**). In contrast, overexpression of all CD97 isoforms led to significant increases in phosphorylated ERK1/ERK2 in most of our PDGCs (**Figure 4C,D**). These results suggested that CD97 activates the MAPK signaling pathway.

**Figure 4:**
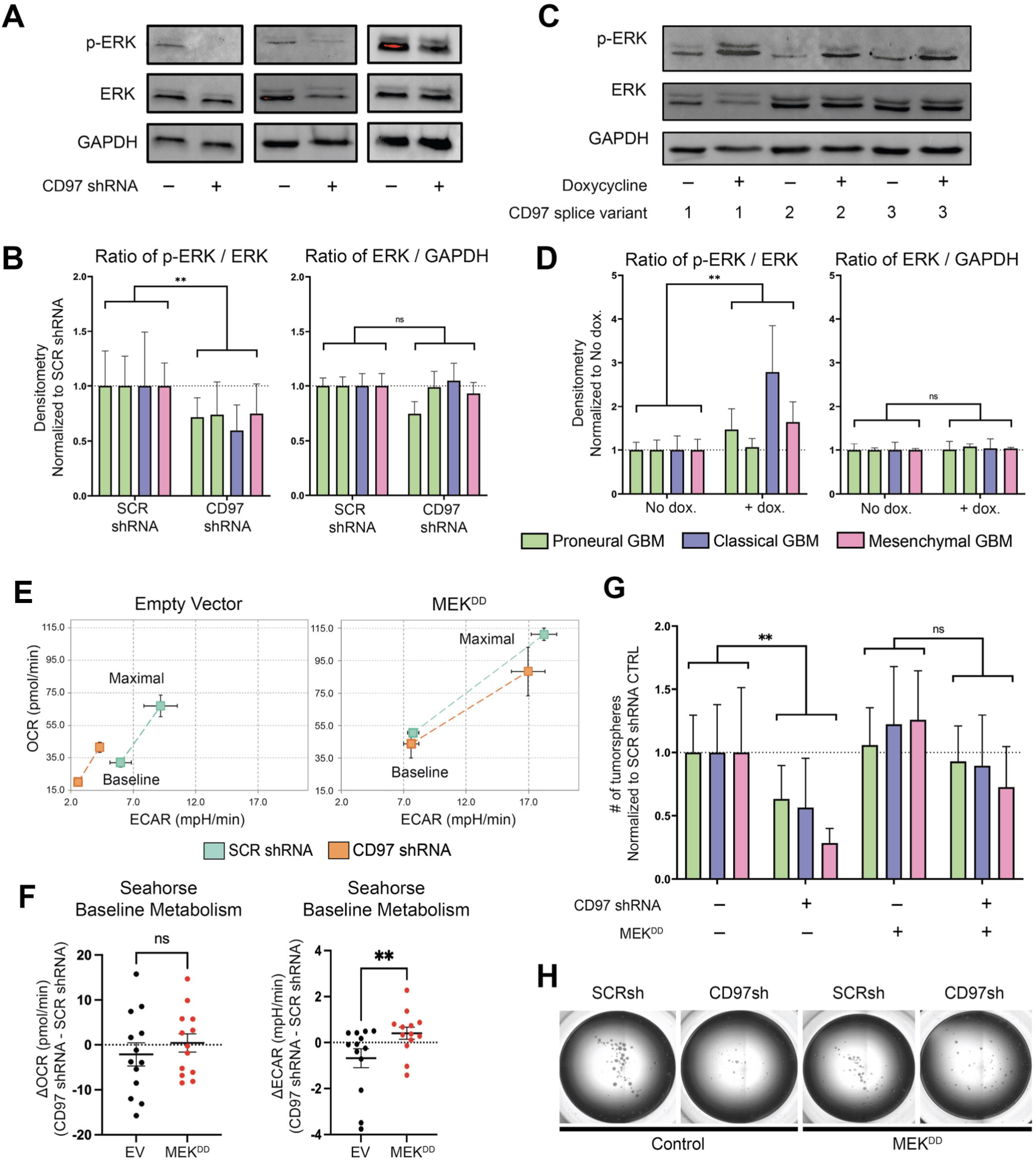
CD97 activates the MAPK signaling pathway to regulate glycolysis and tumorsphere formation. (**A**) Immunoblot for phosphorylated ERK1/ERK2, total ERK1/ERK2, and GAPDH in PDGCs following CD97 knockdown. Red pixels indicate saturation. (**B**) Quantification of densitometry ratios in **Fig. 4A** (p-ERK/ERK: n=5-6 per PDGC; 2-way ANOVA F_1,17_=8.407, p<0.01; ERK/GAPDH: n=5-6 per PDGC; 2-way ANOVA F_1,17_=2.796, p>0.05). (**C**) Immunoblots for phosphorylated ERK1/ERK2, total ERK1/ERK2, and GAPDH in a PDGC following dox-induced overexpression of all three CD97 isoforms. (**D**) Quantification of densitometry ratios in **Fig. 4C** (all isoforms compiled for each PDGC) (p-ERK/ERK: n=4-8 per PDGC; 2-way ANOVA F_1,21_=10.98, p<0.01; ERK/GAPDH: n=4-8; 2-way ANOVA F_1,21_=1.554, p>0.05). (**E**) Seahorse Cell Energy Phenotype graphs showing OCR and ECAR changes before (baseline) and after (maximal) addition of mitochondrial stressors. (**F**) Quantification of the difference in baseline OCR and ECAR between SCR and CD97 knockdown PDGCs after introduction of the MEK^DD^ mutant or an empty vector control [n=13 (all PDGCs compiled); unpaired t-test; ns, p>0.05; **, p<0.01]. (**G**) Bar graph visualizing tumorsphere formation following CD97 knockdown and introduction of the MEK^DD^ mutant (n=8-9 per PDGC; 2-way ANOVA F_3,66_=8.707, p<0.0001; Tukey’s multiple comparisons test: EV SCRsh vs. EV CD97sh: **, p<0.05; MEK^DD^ SCRsh vs. MEK^DD^ CD97sh: ns, p>0.05). (**H**) Representative wells from the assay in **Fig. 4G**. Error bars throughout this figure indicate SEM.

To test whether activation of the MAPK signaling pathway rescues the metabolic and growth phenotypes observed after CD97 knockdown, we stably transfected PDGCs with a phosphomimetic MEK mutant (MEK S218D,S222D; denoted as MEK^DD^ onward), which is expected to constitutively phosphorylate and activate ERK1/ERK2.^50^ We confirmed that the MEK^DD^ mutant increased ERK1/ERK2 activation by performing an immunoblot against phosphorylated ERK (**Supp. Fig. 4A**). Transfection of PDGCs with the MEK^DD^ mutant restored their metabolic profile in Seahorse Cell Energy Phenotype assays following CD97 knockdown (**Figure 4E**). Of particular interest was the finding that MEK^DD^ rescued the CD97 knockdown-imparted defect in ECAR, but not OCR (**Figure 4F**), suggesting the CD97-MAPK signaling axis not only promotes glycolysis but also biases conversion of pyruvate to lactate rather than to acetyl-CoA. Equally importantly, the MEK^DD^ construct also rescued tumorsphere formation after knockdown of CD97 in PDGCs (**Figure 4G,H**). Together, these data suggest that CD97 activates MAPK signaling to promote glycolytic metabolism, thus enabling tumor growth in GBM.

### CD97 activates the MAPK signaling cascade through phosphorylation of its C-terminus and recruitment of β-arrestin

In addition to the MAPK signaling pathway, past research on CD97 has identified other signaling mechanisms, including G protein-mediated Gα_i_ (reduces cyclic-AMP) and Gα_12/13_ (activates RHOA) activation, and PI3K/AKT pathway activation (**Figure 5A**).^27,31,51,52^ We investigated whether CD97 influenced these other signaling pathways in our PDGCs. CD97 overexpression in our PDGCs did not alter canonical G protein-mediated signaling as measured by luciferase reporter assays (**Supp. Fig. 4B**). In addition, CD97 overexpression did not alter Gα_12/13_-mediated activation of RHOA in pulldown assays specific to GTP-bound RHOA (**Supp. Fig. 4C**). Finally, phosphorylation of AKT was unchanged by perturbations in CD97 levels (**Supp. Fig. 4D**). This led us to explore alternative GPCR signaling mechanisms.

**Figure 5:**
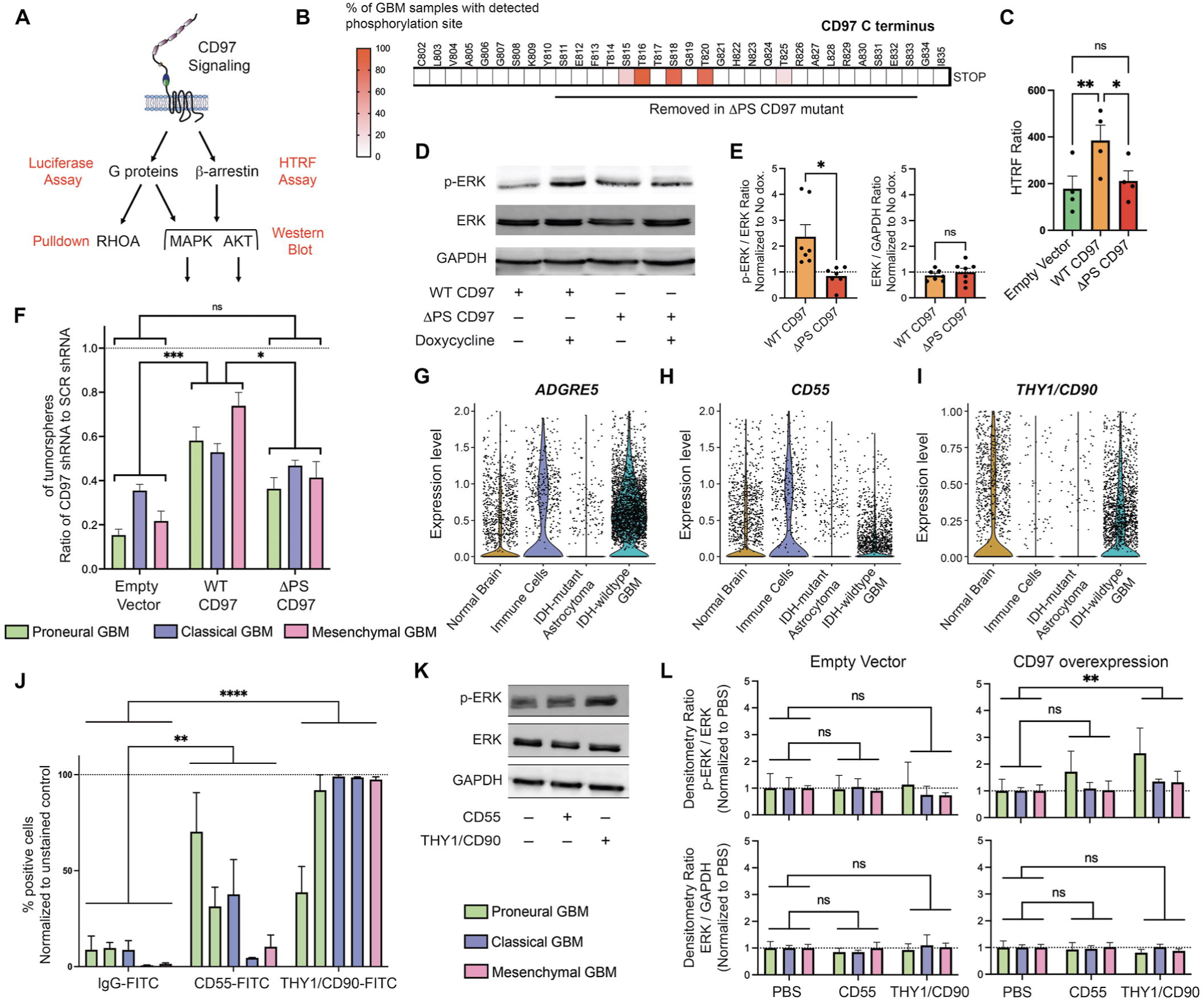
CD97 activates the MAPK signaling cascade through recruitment of *β*-arrestin to its phosphorylated C-terminus and by association with THY1/CD90. (**A**) Schematic of possible CD97 signaling mechanisms tested. Listed in red are the methods for testing. (**B**) Phosphoproteomic data collected from GBM samples identify five phosphorylation sites on the CD97 cytosolic C-terminus. The heatmap depicts the percentage of GBM samples with the detected phosphorylation site. (**C**) HTRF ratios from a β-arrestin recruitment assay performed after overexpression of WT CD97 or the ΔPS mutant. Two PDGCs (n=2 for each) are compiled (n=4 per condition; unpaired t-test; ns, p>0.05; *, p<0.05; **, p<0.01). (**D**) Immunoblot for phosphorylated ERK1/ERK2 after overexpression of WT CD97 or the ΔPS mutant. (**E**) Bar graphs displaying densitometry ratios from **Fig. 5D** (n=7; paired t-test; ns, p>0.05; *, p<0.05). Two PDGCs (n=3-4 for each) are compiled. (**F**) Bar graphs depicting the number of tumorspheres in a tumorsphere formation assay after CD97 knockdown followed by overexpression of shRNA-resistant forms of WT CD97 or the ΔPS mutant. The SCR shRNA groups used for normalization are not shown (n=4 per PDGC; ANOVA F_5,12_=80.79, p<0.0001; Tukey’s multiple comparisons test: EV vs. WT: ***, p<0.001; EV vs. 11PS: ns, p>0.05; WT vs. 11PS: *, p<0.05). (**G-I**) Single-cell RNA/ATAC-seq data from **Fig. 1I** showing expression of *ADGRE5*, *CD55*, and *THY1/CD90*. (**J**) A bar graph depicting the percentage of positive cells measured by flow cytometry after surface staining of PDGCs with FITC-conjugated antibodies against CD55, THY1/CD90, and an IgG control (n=3 for each PDGC; 2-way ANOVA F_2,20_=95.11, p<0.0001; Tukey’s multiple comparisons test: IgG vs. CD55: **, p<0.01; IgG vs. CD90: ****, p<0.00001; CD55 vs. CD90 ****, p<0.01). (**K**) Immunoblot for phosphorylated ERK1/ERK2 from CD97-overexpressing PDGCs plated on laminin or recombinant forms of putative ligands CD55 and THY1/CD90. (**L**) Quantification of densitometry ratios from immunoblots in **Fig. 5K** (n=2-4 per PDGC; Empty Vector p-ERK/ERK: 2-way ANOVA F_2,6_=0.9450, p>0.05; Empty Vector ERK/GAPDH: 2-way ANOVA F_2,6_=1.047, p>0.05; CD97 Overexpression p-ERK/ERK: 2-way ANOVA F_2,16_=12.48, p<0.001; Tukey’s multiple comparisons test: Laminin vs. CD55: ns, p<0.05; Laminin vs. CD90: ***, p<0.001; CD97 Overexpression ERK/GAPDH: 2-way ANOVA F_2,16_=2.257, ns, p>0.05). Error bars throughout this figure indicate SEM.

Many GPCRs are phosphorylated by GPCR kinases (GRKs), a modification which leads to recruitment of β-arrestin.^53^ β-arrestin acts as a scaffold enabling recruitment of components of the MAPK pathway to GPCRs, thereby facilitating subsequent activation of this signaling pathway^54,55^, in addition to its well-described roles in GPCR desensitization and internalization.^63^ To determine whether CD97 undergoes phosphorylation in GBM, we analyzed phosphoproteomic data from GBM in the CPTAC database.^35^ Indeed, we identified five novel serine/threonine phosphorylation sites within the cytosolic C-terminus of CD97 (**Figure 5B**). Three of these phosphorylation sites (T816, S818, and T820) were found in the vast majority of the GBM samples tested and also demonstrated a strong positive correlation in their abundance to one another (**Supp. Fig. 4E,F**). This finding prompted us to test whether CD97 phosphorylation recruits β-arrestin as a potential mechanism for MAPK pathway activation. We generated a CD97 isoform 3 truncation mutant (denoted as ΔPS CD97 onward) lacking amino acids 762-784, which include the five novel and one previously described^52^ phosphorylated residues (**Supp. Fig. 4G**). We verified localization of the ΔPS CD97 mutant to the cell membrane via immunofluorescent staining of the CD97 N-terminus after construct overexpression in HEK293 cells (**Supp. Fig. 4H**). When we performed a fluorescence resonance energy transfer (FRET)-based β-arrestin recruitment assay in GBM cells, we observed enhanced signal upon wildtype (WT) CD97 overexpression, an effect which was no longer observed after overexpression of the ΔPS CD97 construct (**Figure 5C**). Furthermore, overexpression of the ΔPS CD97 mutant was not able to induce phosphorylation of ERK1/ERK2 (**Figure 5D,E**) and failed to rescue tumorsphere formation in PDGCs following CD97 knockdown, as measured by a tumorsphere formation assay (**Figure 5F**). These data suggest that the CD97-MAPK signaling axis requires phosphorylation of the cytosolic C-terminus of CD97 and recruitment of β-arrestin.

### THY1/CD90 is the most likely CD97 ligand in GBM

CD97 is known to interact with several ligands, including CD55, a glycosylphosphatidylinositol (GPI)-linked membrane-tethered protein that regulates the complement cascade in immune cells, and THY1/CD90, another GPI-linked protein found on neurons, immune cells, and activated endothelium.^56-58^ To gain insight into the expression profile of putative CD97 ligands in GBM, we investigated our own, as well as publicly available single-cell RNA-seq data in GBM (**Figure 5G-I**; **Supp. Fig. 5A-F**).^58,59^ While *CD55* expression was largely confined to immune cells, *THY1/CD90* was found in tumor cells, as well as non-neoplastic brain cell lineages (**Figure 5G-I**). That *THY1/CD90* is abundantly expressed in GBM cells, as well as neurons, was corroborated by our initial investigation of transcript levels in our PDGCs and brain cell types (**Figure 1A**). Furthermore, surface staining of PDGCs for CD55 and THY1/CD90 in flow cytometry assays revealed varied expression of CD55, but consistently high levels of THY1/CD90 (**Figure 5J**).

Based on these findings, we then tested whether we could further enhance CD97-activated MAPK signaling in PDGCs by exposure to recombinant forms of CD55 and THY1/CD90. We found that plating CD97-overexpressing PDGCs on wells coated with recombinant THY1/CD90 resulted in increased levels of phosphorylated-ERK1/ERK2 (**Figure 5K)**, an effect that was not observed to the same extent in cells transduced with an empty vector (EV) control (**Figure 5L**). Though CD55 also led to some increases in phosphorylated-ERK1/ERK2, the effect was limited and did not reach statistical significance. Collectively, these data suggest that THY1/CD90 is the most likely activating ligand of CD97 in GBM.

### A CD97-targeting antibody-drug conjugate (ADC) selectively kills PDGCs but not astrocytes or NSCs

Since CD97 exhibits high expression in GBM and is absent from normal brain tissue, we tested whether CD97 could be targeted therapeutically. We generated a human CD97-specific antibody by performing *in vitro* selection of a human synthetic antibody library (**Figure 6A**). We found that the anti-CD97 antibody exhibited significantly increased internalization into PDGCs when compared to an IgG1 LALA isotype control that does not bind to any known target (**Supp. Fig. 6A**). We then conjugated this antibody to monomethyl auristatin F (MMAF), a membrane-impermeant anti-tubulin agent blocking cytoskeletal polymerization and thereby cytotoxic to dividing cells^60^, to generate an anti-CD97 antibody-drug conjugate (ADC). We observed a lower lethal dose (LD_50_) in PDGCs compared to normal brain cells (NHAs and NSCs) in WST8 viability assays (**Figure 6B,C**; **Supp. Fig. 6B-E**). Since the WST8 reagent relies on the presence of cellular metabolites, the assay’s range depends on the metabolic activity of the cells, which varies considerably among PDGCs, NSCs, and relatively non-proliferative NHAs. Therefore, we also stained cells with Hoechst 33342 dye after treatment and took representative images to obtain cell counts (**Figure 6D,E**). From these images, the specificity of the CD97 ADC was even more evident, as only GBM cells decreased in number, while NSC and NHA numbers remained stable. Together, these findings suggest that the *de novo* expression of CD97 in GBM relative to healthy brain can be exploited therapeutically through the use of ADCs.

**Figure 6:**
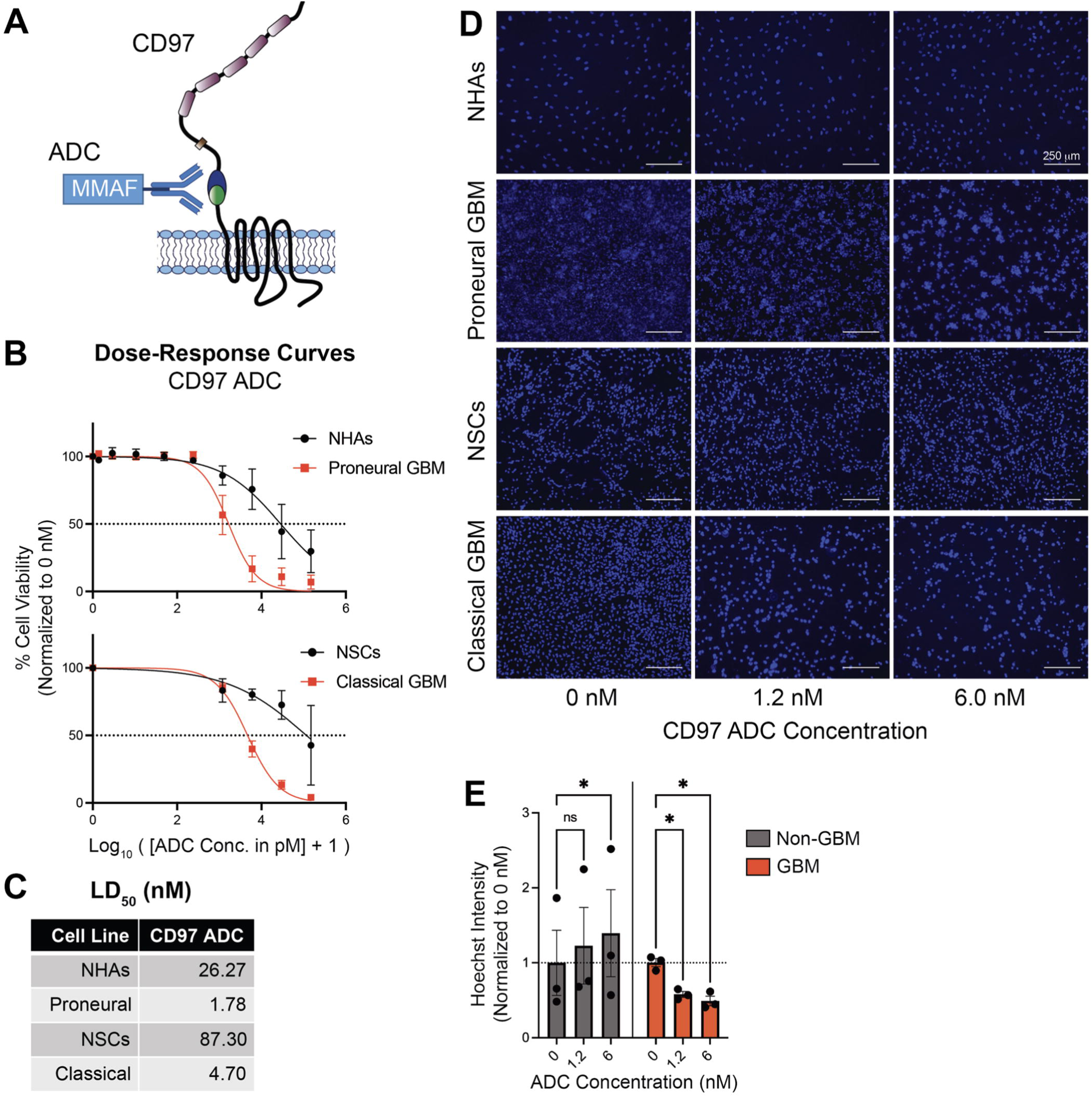
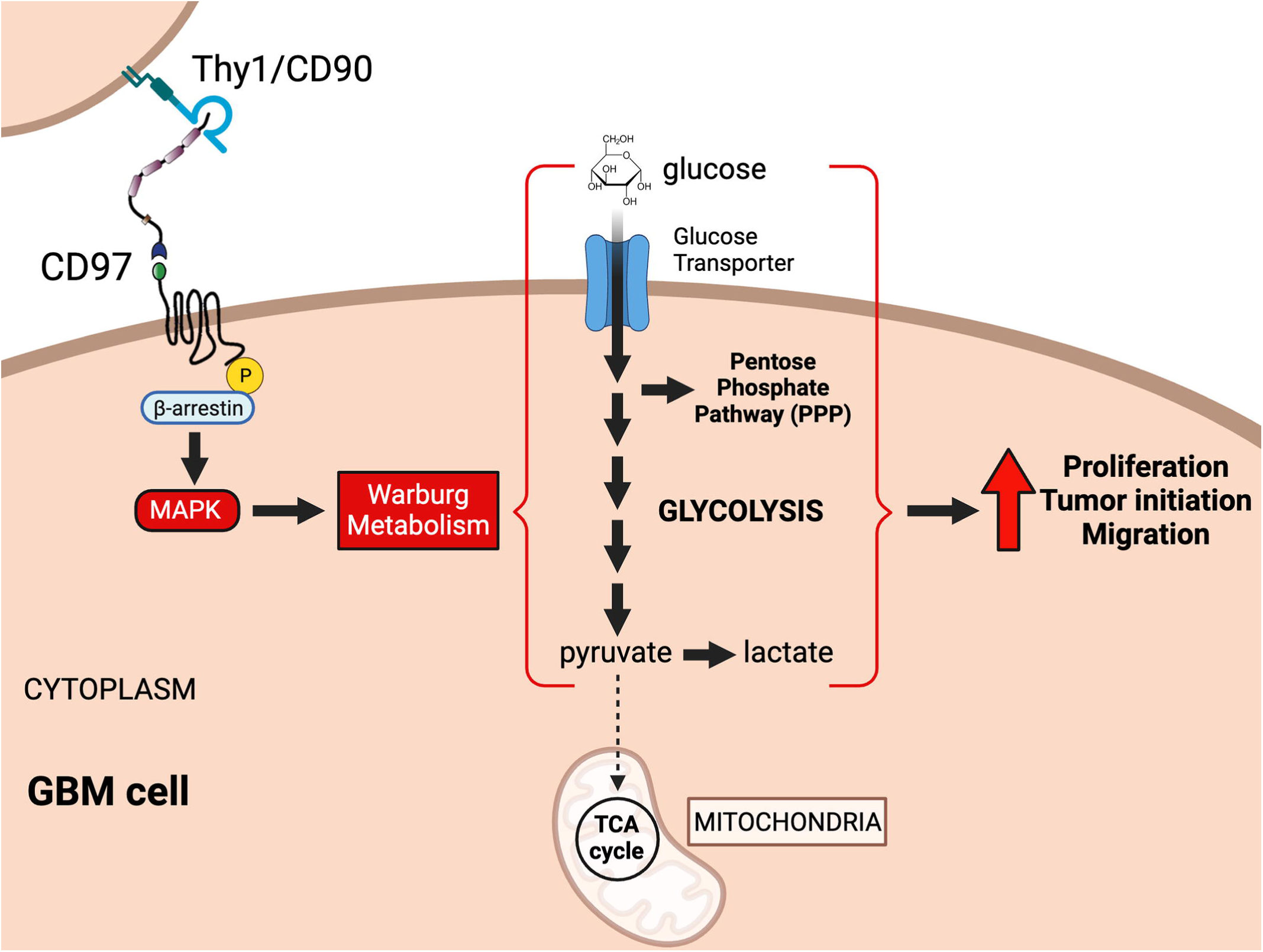
A CD97-targeting antibody-drug conjugate demonstrates enhanced killing of PDGCs compared to NSCs and NHAs. (**A**) Schematic of the MMAF-conjugated ADC binding to the CD97 receptor. (**B**) Dose-response curves generated from WST8 cell viability assays (n=3 biological replicates based on technical triplicates) after treatment of PDGCs, NHAs, and NSCs with the CD97 ADC. (**C**) Table displaying the LD_50_ values (in nM) for the CD97 ADC on all four cell lines as calculated from dose-response curves in **Fig. 6B**. Curves were based on a nonlinear regression fit. (**D**) Representative images of Hoechst 33342-stained cells after six days of treatment with different concentrations of the CD97 ADC. (**E**) Quantification of Hoechst intensity in **Fig. 5D**. Values are normalized to the 0 nM condition. NHAs and NSCs are compiled (non-GBM; black) and the two GBM lines are compiled (GBM; red) to make differences more evident (n=1-2 for each cell line; 2-way ANOVA F_2,8_=12.13, p<0.01; Tukey’s multiple comparisons test: Non-GBM: 0 nM vs. 1.2 nM: ns, p>0.05; 0 nM vs 6 nM: *, p<0.05; GBM: 0 nM vs 1.2 nM: *, p<0.05; nM vs. 6 nM: *, p<0.05). Error bars throughout this figure indicate SEM.

## DISCUSSION

GBM tumors resist multimodal treatment regimens, all but guaranteeing disease recurrence in patients. Identifying new targets for GBM is essential for ameliorating therapy, an already difficult task given GBM’s extensive intertumoral and intratumoral heterogeneity.^3-5,2,19^ Adhesion GPCRs are an understudied group of transmembrane receptors with emerging roles in oncology^10-12^ that show promise as therapeutic targets.^6-9^ Here, we identify the aGPCR CD97 (*ADGRE5*) as a ubiquitously expressed target in GBM, regardless of TCGA transcriptional subtype, thereby overcoming issues of intertumoral heterogeneity. Equally importantly, CD97 is absent from normal non-neoplastic brain tissue, further increasing its potential as a therapeutic target for GBM. In fact, the transcriptional upregulation of *CD97* mRNA in GBM relative to NSCs, the putative cell of origin in glioma, is associated with the highest statistical significance genome-wide (**Figure 1**). Consistent with this expression profile, PDGCs treated with our in-house generated CD97 ADC demonstrated a significantly greater cell death than human astrocytes and NSCs, which lack CD97 expression (**Figure 6**). Collectively, our data suggest that CD97 in GBM is, by virtue of its expression profile, the most appealing treatment target among the aGPCRs, and possibly one of the most targetable cell surface proteins genome-wide. Furthermore, its essential tumorigenic functions in GBM suggest therapeutic targeting cannot only be accomplished via ADC platforms, but that neutralizing antibodies may also be beneficial. A crucial next step in investigating the therapeutic benefits of our CD97 ADC would be to treat orthotopic tumor xenografts *in vivo* with the ADC compound. The treatment schedule and dosage, along with the method of drug administration, whether it be systemic or intracranial/intratumoral, would require optimization. The endogenous expression of CD97 by immune cells and other extracranial tissues warrants concern for peripheral toxicity if the ADC were to be administered systemically in patients. CD97 neutralization with a systemically administered monoclonal antibody, for example, has been found to reduce granulocyte infiltration to sites of inflammation and relieve arthritic activity in mice^24^, corroborating our concern for possible extracranial toxicity of systemically administered CD97 biologics. In order to avoid targeting immune cells or other CD97-expressing tissues, delivery of antibody therapy within the tumor or the brain may be preferred. Novel approaches are being developed to carry out such localized treatments.^61^ Future studies would also be needed in order to test the impact of this CD97-targeting ADC on other CD97-expressing malignancies.

Historically, CD97 was first discovered as a leukocyte receptor and characterized in the context of the immune system.^20-22,62^ Since then, however, CD97 has been implicated in multiple malignancies, including hematologic (leukemia) and solid (esophageal, stomach, colorectal, hepatocellular, pancreatic, thyroid, prostate, ovarian, breast) tumors.^25-28^ CD97 expression in GBM was first demonstrated in 2012, when it was found to confer a migratory phenotype.^33^ More recent work has also implicated CD97 in cellular proliferation, GBM stem cell self-renewal, and tumor metabolism,^29-32^ but the underlying mechanisms have not been elucidated. Here, we have taken a methodical approach to characterize the oncogenic function of CD97 in GBM using PDGCs from all TCGA-defined transcriptional subtypes (proneural, classical, mesenchymal) and advanced genetic, transcriptomic, metabolomic, cellular, and biochemical approaches. Our detailed study not only describes phenotypic contributions of CD97 to cellular proliferation, tumor initiation, GBM stem cell self-renewal, metabolism, and migration, but also presents detailed molecular mechanisms accounting for these phenotypes.

Tumor metabolism, which relies both on the availability of nutrients and their utilization by tumor cells, is a critical determinant of oncogenesis at the interface of tumor cell-intrinsic biology and the microenvironment. GBM, like many other malignancies, manifests a dependency on glycolytic metabolism, rather than tricarboxylic acid (TCA) cycle and oxidative phosphorylation (OXPHOS). The predilection for glycolysis, which occurs in both normoxic conditions and in the severely hypoxic niches of GBM, is known as Warburg metabolism.^39-41,63^ This metabolic adaptation is thought to allow generation of sufficient consumable energy in the form of ATP, but also enable carbon skeleton conservation that is essential for biomass generation in enlarging tumors.^64^ The mechanisms that regulate GBM’s dependency on glycolysis remain incompletely understood. Our data suggest an important role for CD97 in the regulation of GBM cellular metabolism, and glycolysis in particular. *ADGRE5* mRNA transcript levels show a striking positive correlation with transcripts encoding most glycolytic enzymes, and a negative correlation with the expression of most TCA cycle enzymes. Using steady-state metabolomics, we found that CD97 knockdown resulted in a profound depletion of the majority of glycolytic metabolites, and also metabolites within the TCA cycle. Since glycolytic products feed into TCA cycle and mitochondrial metabolism via the conversion of pyruvate to acetyl-CoA, it is not surprising that reduced levels of glycolytic metabolites result in reduced levels of TCA cycle intermediates. Mechanistically, we theorize that CD97 signaling promotes transcription of glycolytic transcripts, including *LDHA* (lactate dehydrogenase A) (**Figure 3C**), which catalyzes conversion of pyruvate to lactate at the expense of the pyruvate to acetyl-CoA reaction that feeds into the TCA cycle. We postulate that the upregulation of certain TCA cycle and OXPHOS transcripts upon CD97 knockdown signifies a compensatory, but ultimately inadequate, adaptation by cells in order to increase ATP production and meet cellular energy needs. More investigation is needed to determine the full mechanism whereby CD97 is impacting rates of glycolysis beyond effects on transcription rates of relevant genes.

The synthesis of our experimental data led us to formulate the hypothesis that CD97’s effects on GBM cellular metabolism and tumor growth are driven by its activation of the MAPK/ERK pathway.^31,51^ This pathway has been implicated in a slew of cellular functions, including cellular metabolism, proliferation, and migration.^65^ From the metabolism point of view, MAPK signaling modulates several enzymes involved in glycolytic and TCA cycle/OXPHOS pathways.^45,66,67^ Mutations that lead to overactivation of the MAPK pathway, such as in *RAS* or *RAF*, are often observed in malignancies and have been linked to their metabolic reliance on glycolysis and the PPP.^49,68^ Pancreatic tumors driven by the oncogenic KRAS G12D mutation, for example, exhibit overactivation of the MAPK pathway, leading to redirection of glycolytic intermediates into the PPP and overall metabolic reprogramming.^49^ Determining the exact mechanism by which CD97-activated MAPK signaling influences cell metabolism and other cell functions is a crucial next step of investigation prompted by this study. For example, whether CD97-MAPK signaling leads to post-translational modifications, such as phosphorylation, of relevant metabolic enzymes remains to be seen.

The impact of CD97 on GBM biology mirrors some of its actions in physiological processes. Within the immune system, CD97 has been shown to be critical for the inflammatory response by mediating cellular adhesion and migration.^20-22,62^ For example, CD97-expressing immune cells have been shown to bind to THY1/CD90-expressing activated endothelial cells^58^, which may represent an important step in leukocyte recruitment and diapedesis at inflamed tissues. Our demonstration of CD97-mediated regulation of cellular metabolism does not necessarily exclude these previously-established functions in inflammation and cellular migration. In fact, it is well-documented that inflammation induces a metabolic shift in immune cells, leading to a greater reliance on glycolysis.^69^ Furthermore, CD97 is expressed in macrophages, neutrophils, and T cells, and is upregulated upon their activation during the immune response, an event accompanied by an increased metabolic reliance on glycolysis.^20,21,48,62,70-73^ The link between metabolism and cell motility is thoroughly documented, but their interplay remains complex.^74-77^ Migratory cells, for example, have been shown to increase glucose uptake by upregulating expression of glucose transporters. This enables localized bursts in glycolysis at sites undergoing cellular contraction needed for cell motility.^74,76^ In this way, CD97’s role in cellular metabolism may complement its previously established role in cellular migration. It is likely that CD97 plays similar functions in activated immune cells and in cancer cells, both of which share predilections for glycolytic metabolism and exhibit extended capacities of migration and proliferation.

Past studies on CD97 have suggested dependencies on multiple signaling mechanisms. A study in prostate cancer cells found that CD97 coupled with Gα_12/13_, ultimately activating the RHOA pathway and promoting cellular migration.^27^ Another study in human retinal pigment epithelium found that overexpression of CD97 activates MAPK signaling, as measured by increased serum response element (SRE)-driven luciferase expression in a luciferase-reporter system.^78^ Interestingly, we did not observe this finding using an SRE-driven luciferase assay in our PDGCs, nor did we detect any other G protein-coupling of CD97 in GBM cells using other luciferase-driven reporter systems (**Supp. Fig. 4B**). We previously found that CD97 activated both MAPK and AKT signaling in leukemic stem cells.^26^ In our PDGCs, we only observed increased levels of phosphorylated ERK1/ERK2, but not phosphorylated AKT (**Supp. Fig. 4D**). A study in colorectal cancer cells demonstrated that CD97 underwent phosphorylation at its C-terminus, which ultimately regulated proper cytoskeletal organization and adhesion.^52^ In a different study, GRK6 was shown to phosphorylate CD97, leading to receptor internalization.^79^ Our analysis of phosphoproteomic data from GBM biospecimens identified robust phosphorylation of novel serine/threonine residues at the cytosolic C-terminus of CD97, a finding that prompted us to investigate the involvement of β-arrestin in CD97-MAPK signaling. β-arrestin is known to act as a scaffold for MAPK components, thereby facilitating pathway activation.^54^ Indeed, we showed β-arrestin recruitment to the phosphorylated C-terminus of CD97. In order to test the role of the phosphorylated C-terminus in CD97 signaling, we generated a ΔPS CD97 mutant lacking the novel phosphorylated serine/threonine residues and found that the truncation abolished β-arrestin recruitment and MAPK activation. A limitation of this experimental approach is that removal of the C-terminus may compromise other aspects of CD97 function, such as G protein-coupling that our assays have failed to detect, or receptor internalization, which may confound interpretation of this result. In the future, we will consider performing site-directed mutagenesis to interrogate each one of the phosphorylated serine/threonine residues individually in relation to CD97 signaling and function in GBM.

Another important aspect of our study is the characterization of CD97-ligand interactions of relevance in GBM. CD97 has a number of known extracellular ligands and binding partners, including the GPI-linked proteins CD55 and THY1/CD90.^58,59^ In contrast to CD55, THY1/CD90, a known neuronal marker, exhibited abundant expression in GBM cells (**Figure 1A,B**; **Figure 5I,J; Supp. Fig. 5D**). When CD97-overexpressing PDGCs were plated on recombinant THY1/CD90 we observed a significantly more pronounced increase in MAPK signaling relative to plating on CD55 (**Figure 5K,L**). Together these data suggest that CD97-activated MAPK signaling most likely relies on the activating ligand THY1/CD90.^57^ Previous studies have implicated THY1/CD90 in GBM pathogenesis and other malignancies.^80,81^ THY1/CD90 is known to form complexes with integrins, which have also been described as CD97 binding partners.^82^ Further insight into the binding interaction between CD97, THY1/CD90, and other putative ligands would help elucidate how crucial this binding is to the phosphorylation of the CD97 C-terminus, recruitment of β-arrestin, and activation of the MAPK signaling pathway. Additionally, the impact of autoproteolytic cleavage, which we observed for both endogenous and exogenous CD97 in PDGCs, on CD97 signaling is a future research aim.

Gliomas are broadly categorized based on their expression of a neomorphic mutation in the IDH enzyme.^5^ Wildtype IDH is found in both the cytoplasm and the mitochondria, where it catalyzes the decarboxylation of isocitrate to α-ketoglutarate (α-KG). The mutant form of IDH, however, generates the oncometabolite 2-hydroxyglutarate (2-HG). In response to 2-HG accumulation, IDH-mutant cells increase levels of OXPHOS and reduce lactate production by silencing *LDHA* expression.^83,84^ This metabolic reprogramming in IDH-mutant glioma lies in stark contrast with that which occurs in IDH-wildtype GBM, where cells are characterized by increased Warburg metabolism. Our single-cell multiomic and ChIP-seq analyses found that *ADGRE5* gene expression is robust in IDH-wildtype GBM, but significantly lower in low-grade IDH-mutant glioma (**Figure 1I**). This finding correlates with the metabolic profiles of the two glioma types and is consistent with our hypothesis that CD97 promotes Warburg metabolism.

In summary, our study identifies CD97 as the aGPCR with the widest differential expression pattern in IDH-wildtype GBM versus normal brain tissue and NSCs, the putative cell-of-origin. We also show that CD97 functions to promote Warburg metabolism via a signaling mechanism that includes phosphorylation of the receptor’s cytosolic C-terminus, recruitment of β-arrestin, and activation of MAPK signaling. Finally, we use a novel CD97-ADC to selectively target GBM cells. Overall, these studies reveal novel insights regarding the role of CD97 in GBM, suggesting that CD97 represents an appealing therapeutic vulnerability in GBM and that its targeting will be impactful in future clinical trials.

## ACKNOWLEDGEMENTS

We thank the Genome Technology Center, the Microscopy Core, the Flow Cytometry Core, the Research Support Service, the Applied Bioinformatics Laboratory, the Center for Biospecimen Research and Development (CBRD) Core, the Vilcek Institute, the Skirball Mouse Facility, and the Metabolomic Laboratory at NYU Grossman School of Medicine. We also thank Drs. Kiyomi Araki and Ben Neel for sharing MEK^DD^ constructs; Dr. Iannis Aifantis and his lab members for use of lab equipment; and Dr. Daniel Orringer for providing one of the surgical specimens. We also thank the Allen Brain Atlas, the Broad Institute, and the Gene Expression Omnibus for their commitment to providing publicly accessible data.

## AUTHOR CONTRIBUTIONS

Conceived and designed the experiments: NR-B, NR, TH, DRJ, RP, AkK, SK, CYP, DGP. Performed and aided with the experiments: NR-B, NR, TH, HD, ChL, CaL, ArK, TR, JDF, GS, JW, ADC, JC, RR, SW, SH, JSO, OM, DZ. Analyzed the data: NR-B, DB, EL, AK-J, AT, DF, FP, DGP. Wrote and edited the manuscript: NR-B, DGP.

## FUNDING

This study was supported by NIH/NINDS R01 NS102665, NIH/NINDS R01 NS124920, NIH/NINDS R21 NS126806, and NYSTEM (NY State Stem Cell Science) IIRP C32595GG to DGP; by R21 CA246457 to TH; and R01 CA251669 to SK and CYP. DGP was also supported by NIH/NIBIB R01 EB028774, NIH/NCI R21CA263402, NIH/NCATS 2UL1TR001445, NYU Grossman School of Medicine, and DFG (German Research Foundation) FOR2149. NR-B was supported by a NYSTEM Institutional training grant (#C322560GG) and an NIH Cell Biology Graduate Student Training Grant (T32GM136542). Core facilities were supported in part by Cancer Center Support Grant P30CA016087 to the Perlmutter Cancer Center of NYU Grossman School of Medicine.

## DECLARATION OF INTERESTS

DGP and NYU Grossman School of Medicine own an EU and Hong Kong patent titled “Method for treating high-grade gliomas” on the use of GPR133 as a treatment target in glioma. SK, TH, AK, CYP, DGP and NYU Grossman School of Medicine have filed a patent application titled “Anti-CD97 antibodies and antibody-drug conjugates”. DGP has received consultant fees from Tocagen, Synaptive Medical, Monteris, Robeaute, Guidepoint, and Advantis. SK was a scientific advisory board member and received consulting fees from Black Diamond Therapeutics; and is a co-founder and holds equity in Aethon Therapeutics and Revalia Bio; has received research funding from Aethon Therapeutics, Argenx BVBA, Black Diamond Therapeutics, and Puretech Health.

## STAR METHODS

### Cell culture

PDGCs were established and maintained as we previously described.^11,37,85^ In brief, fresh operative specimens were obtained from patients undergoing surgery for resection of GBM after informed consent (NYU IRB study 12-01130). Specimens were mechanically minced using surgical blades followed by enzymatic dissociation using Accutase (Cat# AT104, Innovative Cell Technologies). Cells were either long-term maintained in spheroid suspension cultures (tumorspheres) on untreated cell culture dishes or grown as attached cultures on dishes pretreated with poly-L-ornithine (PLO; Cat# P4957, Sigma) and laminin (Cat# 23017015, Thermo Fisher). The GBM growth medium consisted of Neurobasal medium (Cat# 21103049, Gibco) supplemented with N2 (Cat# 17-502-049, Gibco), B27 (Cat# 12587010, Gibco), nonessential amino acids (Cat# 11140050, Gibco), and GlutaMax (Cat# 35050061, Gibco). Twenty ng/ml recombinant basic Fibroblast Growth Factor (bFGF; Cat# 233-FB-01M, R&D) and 20 ng/ml Epidermal Growth Factor (EGF; Cat# 236-EG-01M, R&D) were added to the medium every other day. Parental tumors of these patient-derived cultures underwent DNA methylation, mutational and copy number variation profiling using previously described assays.^37^ All tumors used for PDGCs had a wildtype IDH genetic background.

Human embryonic stem cell (WA09) derived NSCs were established and maintained as we previously described in detail.^16^ HEK293T (Cat# 632180, Takara) cells were cultured in Dulbecco’s modified Eagle’s medium (DMEM; Cat# 11965-118, Gibco) supplemented with 10% fetal bovine serum (FBS; Cat# PS-FB2, Peak Serum) and sodium pyruvate (Cat# 11360070, Gibco). NHAs (Cat# CC-2565, Lonza) were cultured in PLO/laminin-coated tissue culture plates in DMEM media with 10% FBS and N2 supplement. All cells were cultured in humidified cell culture incubators at 37°C balanced with 5% CO_2_ and at 21% O_2_.

### Flow cytometry surface staining

Cells were enzymatically dissociated using Accutase and 5×10^5^ cells were pelleted at 1000 rpm for 5 minutes. Cells were washed in Dulbecco’s phosphate-buffered saline (DPBS; Cat# 14190-144, Gibco) before resuspension in fluorescence-activated cell sorting (FACS) buffer [0.5% BSA (Cat# A3733-100G, Sigma) in DPBS with 2 mM EDTA (Cat#AM9260G, Thermo)] with diluted primary antibody for 30 minutes at 4°C. Cells were washed in FACS buffer before a final resuspension in FACS buffer and subsequent transfer to a 5 mL round bottom polysterene test tube through a cell strainer (Cat# 352235, Falcon). Samples were run on a BD LSRFortessa (BD Biosciences). APC- and FITC-conjugated primary antibodies were used for staining against CD97, CD55, and THY1/CD90 based on their commercial availability and high detectability.

### Immunofluorescent staining of tissue

GBM tumors were collected as surgical specimens and normal brain tissue was collected from patients *post mortem*. Tissues were washed for paraffin-embedding and sectioning by the Center for Biospecimen Research & Development (CBRD) core at NYU. Slides were deparaffinized and rehydrated by submersion into xylene (15 minutes) (Cat# 1330-20-7, Crystalgen), 100% ethanol (3 minutes) (Cat# UN1170, Fisher bioreagents), 95% ethanol (3 minutes) (Cat# E7148-500ML, Sigma-Aldrich), and 70% ethanol (5 minutes) (Cat# 2401, Decon Laboratories). The slides were washed three times in DPBS with 0.1% Tween-20 (PBS-T; Cat# H5151, Promega). Antigen retrieval was performed by submerging the slides in citrate buffer [2.94 g sodium citrate tribasic dihydrate (Cat# S4641-500G, Sigma-Aldrich), 0.5 mL Tween-20, and 1000 mL ddH_2_O] and microwaving for 10 minutes at 800 W followed by 10 minutes at 200 W. The slides were then allowed to cool slowly at room temperature and were washed three times with PBS-T. Slides were blocked for 30 minutes using 50 μL normal goat serum (Cat# ab7481, Abcam) in 1 mL DPBS. Slides were then stained with primary antibody solution at 4°C overnight. Slides were then washed three times with PBS-T and stained with secondary antibody for one hour at room temperature in the dark. Slides were washed, stained with 1:2000 Hoechst 33342 dye (Cat# H3570, Life Technologies) or 100X DAPI (Cat# D8417-1MG, Sigma) in DPBS, covered in ProLong gold antifade reagent (Cat# P36934, Thermo), and mounted with a coverslip (Cat# 10813, Ibidi) for imaging.

### Plasmids and molecular cloning

All overexpression plasmids were based on the lentiviral vector pCW57-RFP-P2A-MCS (Plasmid #78933, Addgene). ΔPS CD97 was generated using primers against the CD97 isoform 3 expression vector and was assembled using Gibson Assembly. Short hairpin RNA design and cloning into the pRSI9-U6-(sh)-UbiC-TagRFP-2A-Puro (Plasmid #28289, Addgene) shRNA expression vector have been described previously.^26^ Guide RNAs were designed using Benchling, selected based on off-target scores, and cloned into the LentiCRISPRV2-mCherry (Plasmid# 99154, Addgene) backbone. The MEK^DD^ construct was in a pBabe-neomycin (Plasmid #1767, Addgene) expression vector.

### Lentiviral infection and transfection

Cells were transduced using lentivirus as described previously.^86^ In short, lentivirus was produced by co-transfecting HEK293T cells with expression plasmids of interest and packaging plasmids psPax2 and pMD2.G. Lentivirus was collected from the cell culture supernatant 24, 48, and 72 hours after transfection and concentrated using the Lenti-X concentrator (Cat# 631231, Contech Takara). For lentiviral transduction, GBM, or NHA medium was supplemented with 4 μg/ml protamine sulfate and cells were treated with viral particles at a multiplicity of infection (MOI) of three. Infected cells were isolated by FACS with the SH800Z sorter (Sony Biotechnology) or by puromycin selection (5 μg/mL). Doxycycline induction was done by adding 1 μg/mL doxycycline to the medium. HEK293T cells were transfected with plasmid DNA using Lipofectamine 2000 (Cat# 11668-019, Invitrogen), following the manufacturer’s protocol.

### Immunoblot staining & sample deglycosylation

Cell medium was aspirated and cells were washed with DPBS (Cat# 14190-250, Gibco). RIPA buffer (Cat# 89901, Thermo) with Protease/Phosphatase inhibitor (Cat# 88669, Fisher) was added directly to cells, which were incubated on ice for 15 minutes. A cell scraper was used to collect the whole cell lysate, which was transferred to a 1.5 mL tube (Cat# C2170, Thomas Scientific). Samples were sonicated (ON: 15s, OFF: 60s, 8 cycles, High setting, 4°C) using a Bioruptor300 (Diagrenode). Samples were spun at 15000 rcf for 10 minutes at 4°C and the supernatant was transferred to a fresh tube. If samples were deglycosylated, the Protein Deglycosylation Mix II (Cat# P6044, NEB) was used according to the manufacturer’s protocol and samples were incubated at 25°C for 30 minutes and at 37°C for 16 hours. Bovine serum albumin (BSA) protein standards and a detergent compatible (DC) Protein Assay Kit (Cat# 5000112, BioRad) were used to measure protein concentrations of the whole cell lysates according to the manufacturer’s protocol. Up to 40 μg whole cell lysate was mixed in 4X Laemmli Buffer (Cat# 1610747, BioRad) and run in a 12% Mini-PROTEAN TGX Precast Protein Gel (Cat# 4561044, BioRad) for 1 hour at 120 V. Protein gels were then transferred to a PVDF membrane (Cat# 1620177, BioRad) for 2 hours at 90 V at 4°C. Membranes were blocked [2% BSA in tris-buffered saline with 0.1% Tween-20 (TBS-T)] for 1 hour at room temperature. Membranes were stained with primary antibody overnight at 4°C. Membranes were then washed three times in TBS-T for 5 minutes before incubation with a secondary antibody for 1 hour at room temperature in the dark. The membrane was finally washed three times in TBS-T for 5 minutes and imaged using an iBrightFL1000 (Invitrogen). Bands for non-phosphorylated species were imaged using fluorescent secondary antibodies and by measuring fluorescent signal. Bands for phosphorylated species were imaged using an HRP-conjugated antibody and by measuring chemiluminescent signal using appropriate West Pico PLUS chemiluminescent substrates (Cat# 34577, Thermo) according to the manufacturer’s protocol.

### Hoechst staining

Cells were resuspended (7 days post infection) in DPBS with 1:1000 Hoechst 33342 dye and incubated at 37°C for 30 minutes. Cells were spun down and resuspended in FACS buffer and transferred to tubes for flow cytometry.

### Annexin-V Staining

One million cells were pelleted (7 days post infection) and washed once in DPBS and then washed once in 1X Binding Buffer (Cat# BMS500BB, eBioscience) in ddH_2_O. Cells were resuspended in 100 μL 1X Binding Buffer with 5 μl Annexin-V eFluor450 (Cat# 88-8006-74, eBioscience) and were incubated for 15 minutes in the dark at room temperature. Three hundred μL 1X Binding Buffer was added to the cells and they were spun down and resuspended in 200 μL 1X Binding Buffer with 100X DAPI solution. After a 10-minute incubation in the dark at room temperature, cells were resuspended in FACS buffer and transferred to a tube for flow cytometry. Cells that were incubated at 56°C for 20 minutes were used a positive control for non-viable cells. Cells that were either Annexin-V positive (PE-Cy7) and/or DAPI (Pacific Blue) positive were considered non-viable.

### Transwell migration assay

PDGCs were dissociated using Accutase and counted (4 days post infection). The cells were resuspended to a concentration of 50,000 cells/100 μL in serum-free Neurobasal media. Meanwhile a 1:30 Matrigel (Cat# 354277, Thermo) to Neurobasal medium solution was made and added to the top well of a transwell permeable support 24-well plate (Cat# 3422, costar) and incubated at room temperature for two hours. The Matrigel solution was aspirated and the well was washed once with DPBS. One hundred μL of cells were added to the top well. The bottom well was filled with 600 μL of Neurobasal medium with 10% FBS. The plate was incubated at 37°C for 24 hours. After 24 hours, the media was aspirated and replaced with medium containing 1:1000 Hoechst 33342 dye and was incubated for 10 minutes at 37°C. The media was then aspirated and replaced with DPBS. Three fluorescent images were taken of the cells on the membrane (total cells). The cells in the upper portion of the well were gently removed using a cotton swab. Three fluorescent images were taken of the remaining cells in the lower portion of the well (migrated cells). The fluorescent intensity was measured and the fraction of migrated cells to total cells was calculated.

### Tumorsphere formation assay

Cells were enzymatically dissociated using Accutase and counted using Countess II (Invitrogen). Cells were spun down at 1000 rpm for 5 minutes and brought to a concentration of 100 - 500 cells / 50 μL. One row (12 wells) of a 96-well plate (Cat# 7200656, Fisher) were filled with 50 μL of the cell suspension. Cells were fed every two days with EGF and bFGF supplements and were given 14 days to form tumorspheres. Ninety-six-well plates were then scanned using an Evos Cell Imaging System (Evos) and tumorspheres were counted using ImageJ software.

### Limiting dilution assays

Short hairpin RNA transduced PDGCs were dissociated using Accutase and counted. Cells were plated in 24 wells of a 96-well plate at four separate concentrations: 1000 cells/well, 300 cells/well, 30 cell/well, and 3 cells/well. Cells were then fed every two days and given 14 days to grow. The number of completely empty wells were recorded. Clonogenic frequencies were calculated using ELDA software (https://bioinf.wehi.edu.au/software/elda/). For plotting, the log_10_ of each frequency was taken.

### Cell competition assay

GBM cells were lentivirally infected with a LentiCRISPRv2 gRNA:Cas9:mCherry construct with an MOI of 3. Cells were then enzymatically dissociated using Accutase into single cells and counted. We mixed 100,000 infected, mCherry+ cells with 100,000 uninfected cells. Flow cytometry [BD LSRFortessa (BD Biosciences)] was used to measure the initial proportion of mCherry+ cells. Every four days, tumorspheres were enzymatically dissociated, the proportion of mCherry+ cells was measured via flow cytometry, and 100,000 cells were replated. This was done until day 20.

### Luciferase Assay

PDGCs lentivirally infected with a dox-inducible CD97-overexpression or empty vector were transfected with luciferase signaling-reporter plasmids: cAMP response element–Luciferase (Cat# E8471, Promega), SRE-Luciferase (Cat# E1340, Promega), SRF-RE-Luciferase (Cat# E1350, Promega), and NFAT-RE-Luciferase (Cat# E8481, Promega). Twenty-four hours after transfection, cells were reseeded in black 96-well plates at a density of 75,000 cells per well with medium containing doxycycline (or dox-free medium). Forty-eight hours after transfection, cells were lysed and luciferase activity was detected using the Bright-Glo Luciferase assay system (Cat# E2650, CisBio) and a BioTek Synergy H1 microplate reader according to the manufacturer’s protocol.

### *In vivo* GBM xenografts

Mice were housed within NYU Langone Medical Center’s Animal Facilities. All animal procedures were performed according to an IACUC-approved protocol. Orthotopic intracranial xenografts have been described in detail previously.^87^ In short, equal numbers of male and female immunodeficient NSG (NOD.Cg-*Prkdc^scid^ Il2rg^tm1Wjl^*/SzJ) mice (6-8 weeks of age) were anesthetized with intraperitoneal injection of ketamine/xylazine (10 mg/kg and 100 mg/kg, respectively). A midline skin incision was made and a small hole was drilled through the skull 2 mm off the midline and 2 mm anterior to the coronal suture. Mice were then stereotactically injected with 2.5 x 10^5^ GBM cells lentivirally infected with a luciferase-containing plasmid. The skin incision was sutured and animals were closely monitored during the recovery period. Mice were weighed every two weeks and sacrificed once they had lost 20% of their maximum body weight or according to ethical guidelines.

### IVIS imaging of *in vivo* GBM xenografts

*In vivo* GBM xenografts were monitored using an IVIS Lumina XR (PerkinElmer) as described previously.^88^ First, mice were weighed and injected intraperitoneally with 10 μL/g body weight Luciferin substrate solution [D-Luciferin Potassium Salt (LUCK-300, Gold Biotechnology) diluted in DPBS to a final concentration of 20 mg/mL]. Mice were anesthetized using isoflurane and inserted into the IVIS imaging system. Thirteen minutes after Luciferin injection, mice were imaged at a 150 second exposure time. Living Image software (PerkinElmer) was used to quantify the “Radiance (Photons)” within the selected ROI.

### Bulk RNA-sequencing

Three replicates of a PDGC (knockdown, scrambled, overexpression, and empty vector) and one replicate each of three PDGCs (knockdown and scrambled) were plated adherently in a 6-well plate. Two replicates of NSCs were also plated adherently on a 6-well plate. RNA was extracted using a RNeasy Mini Kit (Cat# 74104, Qiagen) according to the manufacturer’s protocol. Samples were submitted to the NYU Langone Genome Technology Core (GTC) for sequencing on an Illumina NovaSeq 6000 flow cell. FastQ files were aligned to the reference genome (hg19) using the splice aware aligner STAR (v2.5.0c), and the read counts were generated. DESeq2 was used for normalization, differential gene expression analysis, and to generate volcano plots. Log-processed RPKM values were used to generate a heatmap of aGPCR expression in GBM cells using Seq-N-Slide (rna-star-groups-dge) software. Peaks were visualized at the *ADGRE5* locus (chr19:14488844-14534630; hg19 reference genome) using the WashU Epigenome Browser (http://epigenomegateway.wustl.edu/browser/) and Integrative Genomics Viewer (IGV).

### Single-Cell 10X Multiome (ATAC + RNA) Sequencing

Fresh IDH-mutant astrocytoma and IDH-wildtype GBM surgical specimens were immediately flash frozen in liquid nitrogen. Tissue was homogenized using a pestle (Cat#12-141-367; Thermo). The sample was then processed as described in the “10X Genomics Nuclei Isolation from Complex Tissues for Single Cell Multiome ATAC + GEX Sequencing Demonstrated Protocol CG000375 Rev B” and nuclei were processed for sequencing on an Illumina NovaSeq 6000 flow cell. Data were processed, normalized, integrated for the two specimens, and clustered using Seurat and Signac packages based on RNA and ATAC peak data in R. Only cells with 500-5000 features and with a percentage of mitochondrial genes less than 15% were included in the analysis. Clusters were identified based on the expression of relevant markers (neuronal, oligodendrocytic, astrocytic, and immune) and based on upregulated enriched pathways determined by GO. Oligodendrocytic and neuroglial clusters were compiled together as “Normal brain cells”. Immune populations from the two datasets were clustered together as “Immune cells”. IDH-mutant astrocytoma and IDH-wildtype GBM clusters were identified by their high astrocytic expression profile along with high expression of other oncogenic markers (*EGFR*, *TIMP3*, *SOX2*, *TUBA1A*, *MYC*, *HIF1A*, *FN1*, *HK2*, *TMOD1*, and *IGFBP5*). ATAC-seq data was visualized at the *ADGRE5* locus (chr19:14380000-14420000; hg38 reference genome) for the clusters using the Signac package in R.

### Pathway enrichment analysis & correlation matrix analysis

Differentially expressed downregulated genes were plugged into GO PANTHER software (http://pantherdb.org/). The top 10 depleted pathways upon CD97 knockdown were plotted based on log_10_(fold enrichment) values. GSEA software (https://www.gsea-msigdb.org/gsea/index.jsp) was used to generate GSEA plots from gene count data. DGCA in R was used to generate a correlation matrix based on RNA-seq data after CD97 knockdown or overexpression in PDGC replicates. Genes designated as glycolytic stem from GO:0061621 along with select glucose transporters. Genes designated as TCA-cycle/OXPHOS include those from GO:0006099 and all cytochrome oxidase subunits. Positive integers designate positive correlation in gene expression patterns, while negative integers designate negative correlation in gene expression patterns.

### Steady-state metabolomics

Three replicates of two PDGCs (knockdown and scrambled) were plated non-adherently on 6-well plates. After two days, all cells were collected into a 1.5 mL tube and spun at 3000 rcf for 1 minute at 4°C. The supernatant was aspirated and cells were washed in 1 mL of DPBS and immediately pelleted again. Cell pellets were flash-frozen and submitted to the Metabolomics Laboratory for hybrid metabolomics analysis using liquid chromatography/mass spectrometry (LC-MS/MS). A more detailed description of the methods performed for metabolite extraction, LC-MS/MS hybrid metabolomics, and relative metabolite quantification is provided in **Supplementary Methods**. One replicate was excluded for technical reasons.

### Flux metabolomics

Three replicates of a PDGC were plated on 6-well plates (knockdown and scrambled). Cells were given fresh Neurobasal-A medium, no D-glucose, no sodium pyruvate (Cat# A2477501, Thermo) with 25 mM [U-^13^C_6_]-glucose (Cat# 389374-10.00G, Sigma). Triplicates were collected and spun at 3000 rcf for 1 minute at 4°C at three separate timepoints (5, 30, & 120 minutes). Cell pellets were washed with 1 mL DPBS and flash-frozen. Tubes were submitted to the Metabolomics Laboratory for hybrid metabolomics analysis using LC-MS/MS. A more detailed description of the methods performed for metabolite extraction, LC-MS/MS with hybrid metabolomics, and relative quantification is provided in **Supplementary Methods**.

### Lactate assay

Lentivirally infected GBM cells were plated on PLO/laminin treated 6-well plates in 1 mL of GBM medium and given 24 hours to grow. A lactate assay cocktail was made [50% Glycine/Hydrazine 0.6M solution (Cat# G5418, Sigma), 1% 240 mM NAD+ (Cat# N0632, Sigma), 2 μL/mL Lactate Dehydrogenase (5U/uL) (Cat# 10127230001, Roche), 49% sterile ddH_2_0]. A series of lactate standards ranging from 10 mM to 0 mM were made (Cat# AC18987-0050, Fisher). Two hundred μL of the lactate cocktail was added to the wells of a 96-well plate. Five μL of each standard or sample was added in triplicates to the wells containing the lactate cocktail. The solution was mixed and incubated for 1 hour at 37°C. The solution was mixed again and the absorbance at 340 nm was read using a Synergy H1 Plate Reader (BioTek).

### Seahorse Cell Energy Phenotype assay

Seahorse XFe24 Analyzer (Agilent) and the Seahorse XF Cell Energy Phenotype Test Kit (Cat# 103325-100, Agilent) were used to perform assays according to the manufacturer’s protocol. In brief, a Seahorse cell culture plate was treated with PLO/laminin and 10,000 dissociated GBM cells were plated per well, with five technical replicates per condition accompanied with a blank control well. Cells were incubated overnight at 37°C in a CO_2_ incubator to allow for attachment. Meanwhile, a Seahorse Utility plate/sensor cartridge was hydrated and calibrated using Calibration buffer (Cat# 100840-000, Agilent) in a non-CO_2_ incubator at 37°C overnight. Cell medium was aspirated and replaced with 500 μL Assay medium [Agilent Base Medium (Cat# 102353-100, Agilent), 1 mM pyruvate (Cat# 103578-100, Agilent), 2 mM glutamine (Cat# 103579-100, Agilent), 10 mM glucose (Cat# 103577-100, Agilent)] and was incubated for 1 hour in a non-CO_2_ incubator at 37°C. The mitochondrial stressors carbonyl cyanide p-(tri-fluoromethoxy)phenyl-hydrazone (FCCP) and oligomycin were prepared to a final molar concentration of 1.25 μM and 10 μM, respectively, and were added to Port A of the sensor cartridge. The Seahorse cell plate and sensor cartridge were run on a Seahorse XF Cell Energy Phenotype Assay program using Agilent Wave software. Both OCR and ECAR were measured at three baseline timepoints and six timepoints after the addition of the mitochondrial stressors.

### MEK mutant transfection & selection

GBM cells were transfected with the neomycin-resistant MEK^DD^ construct or a neomycin-resistant empty vector control using Lipofectamine Stem Transfection Reagent (Cat# STEM00008, Thermo) overnight. Cells were treated for three days with media containing 1 mg/mL G418 sulfate (Cat# 61-234-RG, Corning) dissolved in DPBS. Cells were given 7-14 days to expand in low dose (100 μg/mL) G418 medium before they were used for specified experiments.

### RHOA-pulldown Assay

The RhoA Pull-down Activation Assay Biochem Kit (Cat# BK036, Cytoskeleton) was ordered and the manufacturer’s protocol was followed. Five hundred μg of cell lysate was used and 15 μL beads.

### HTRF-based β-arrestin recruitment assay

GBM cells overexpressing wildtype CD97, the ΔPS mutant, or an empty vector control were plated (50,000 cells/well) on a PLO/laminin coated white 96-well plate. A homogenous time-resolved fluorescence (HTRF)-based β-arrestin2 recruitment kit was used and followed according to the manufacturer’s protocol (Cat# 62BDBAR2PEB, cisbio). HTRF values were measured using a FlexStation 3 Multi-Mode Microplate Reader (Molecular Devices).

### Ligand-based immunoblots

Six-well plates were coated with PLO/laminin. Wells were then additionally coated with recombinant human CD55 (Cat# 2009-CD-050, R&D) or recombinant human THY1/CD90 (Cat# 16897-HCCH, Sino Biological) at a concentration of 24 μg/mL overnight. GBM cells overexpressing CD97 or an empty vector control were seeded (∼1×10^6^ cells/well) on the plate. Whole cell lysates were collected for immunoblotting 24 hours after seeding.

### Generation of antibody-drug conjugate (ADC) against CD97

The gene encoding the ectodomain of CD97 isoform 1 was synthesized (Integrated DNA Technologies) and cloned into the mammalian expression vector pBCAG.^89,90^ The expi293F cells (Thermo Fisher) were transfected with the vector using the ExpiFectamine 293 Transfection Kit (Thermo Fisher) according to the manufacturer’s protocol. The CD97 protein was purified from the cell culture supernatant by immobilized metal affinity chromatography (IMAC) followed by size exclusion chromatography. The sorting of a synthetic human antibody (sAb) library and identification of anti-CD97 sAb by phage ELISA were performed essentially as described previously.^91,92^ The gene encoding the VH domain of the antibody was cloned into a modified version of pFUSEss-CHIg-hG1 (InvivoGen) harboring the LALA-PG (L234A, L235A, P329G) mutations that abrogate Fc receptor binding,^93^ and the gene encoding the VL domain into pFUSEss-CLIg-hk (InvivoGen). The antibody in this IgG1-LALA-PG format was produced using the expiCHO expression system according to the manufacturer’s protocol (Thermo Fisher), and purified with Protein A affinity chromatography. The drug conjugation reaction was performed as described previously.^94^ Briefly, the interchain disulfide bonds of the antibody were cleaved with DTT. A 9.5-fold molar excess of MC-Val-Cit-PAB-MMAF (BOC Science) was added to the reduced antibody, and after 1 hour of incubation, the reaction was quenched by adding excess cysteine. The excess drug and cysteine were removed with the Zeba spin desalting column (Thermo Fisher). Approximately 6-7 drug molecules were conjugated per antibody.

### Antibody internalization assay

Two PDGCs were seeded in 96-well plates in triplicates. Antibodies were conjugated with the pHAb amine reactive dye (Cat# G9841, Promega) according to the manufacturer’s protocol. Dye-conjugated antibodies were purified using the Capturem Protein G miniprep columns (Cat# 635725, Takara). Purified antibodies were dialyzed into DPBS overnight. The concentration of both the anti-CD97 antibody and the isotype control used for the assay was 30 μg/ml and the cells were incubated for 20 hours. The cells were then harvested and analyzed by flow cytometry to determine internalization. The pH sensitive pHAb dye that is conjugated to each antibody has low or no fluorescence (PE channel) at pH>7. Upon internalization, the dye becomes fluorescent at an acidic pH present in early endosomes and lysosomes. The increase in mean fluorescence intensity was measured via the PE channel.

### Antibody-drug conjugate dose-response curves

Ninety-six-well plates were coated with PLO/laminin. We then plated 10^4^ cells per well (100 μL per well; each condition in triplicates) and allowed them to adhere overnight. Serial dilutions of ADC, MMAF, and MMAE were prepared in appropriate cell medium to give the following final concentrations: 150 nM, 30 nM, 6 nM, 1.2 nM, 240 pM, 50 pM, 10 pM, 3 pM, 1.4 pM, and 0. Ten μL from dilutions were added to wells. On the seventh day, cells were stained using 1:2000 Hoechst 33342 dye (Cat# H3570, Life technologies) and representative images were taken. Ten μL of WST8 (Cat# ab228554, Abcam) was added to each well and the absorbance at 460 nm was measured using a Synergy H1 Plate Reader (BioTek) after a 3-hour incubation at 37°C. Dose response curves and respective LD_50_ curves were generated using GraphPad Prism software (version 8.4.3). Curves were based on a nonlinear regression fit.

### Databases used and data availability

The Allen Brain Map Human M1 10X Transcriptomics Explorer database was used to visualize expression levels in normal human brain cells. Proteomic data for normal brain was collected and annotated as described in Perna et al.^34^ Publicly available H3K27ac ChIP-seq from the Gene Expression Omnibus (GEO) was displayed via the Integrative Genomics Viewer (IGV) (https://software.broadinstitute.org/software/igv/). Peaks associated around the *ADGRE5* locus (chr19:14488844-14534630; hg19 reference genome) are shown for all specimens [IDH-wildtype GBM: GSM1824800, GSM1824811; IDH-mutant anaplastic astrocytoma: GSM1824813, GSM1824808 (anaplastic astrocytoma)]. GBM phosphoproteomic and proteomic data were from the GBM datasets of the Clinical Proteomic Tumor Analysis Consortium (CPTAC) included in Wang et al. analyzing multi-omic profiles from 99 treatment-naïve GBM specimens.^35^ Publicly available single-cell RNA-seq data of adult and pediatric glioblastoma from the Broad Institute Single Cell Portal (GSE131928) were used to look at expression of *ADGRE5* and its putative ligands *THY1/CD90* and *CD55*. Bulk RNA-sequencing and single cell RNA/ATAC-sequencing data have been deposited at GEO under GSE230393 and GSE230389 accession numbers, respectively.

### Statistical analysis

All experiments were performed in biological replicates of at least three repeats unless otherwise specified. Statistical analysis was performed using GraphPad Prism (version 8.4.3). Summary statistics are represented as mean ± standard error of the mean (SEM) unless otherwise indicated. Statistical significance was calculated using either Students t-test, logrank test (for Kaplan-Meier survival curves), 1-way analysis of variance (ANOVA), or 2-way ANOVA, with Tukey’s or Sidak’s *post hoc* test for multiple comparisons. P values <0.05 were considered statistically significant (*, p<0.05; **, p<0.01; ***, p<0.001; ****, p<0.0001).

