## Supplementary figures and methods for "The expression profile and tumorigenic mechanisms of CD97 (ADGRE5) in glioblastoma render it a targetable vulnerability"

### SUPPLEMENTARY FIGURE LEGENDS

**Supplementary Figure 1: Additional expression data.** (**A**) Representative histogram data from **Fig. 1C**, showing CD97:APC staining versus an IgG:APC control. (**B**) Dot plot displaying gene expression profiles of the single-cell clusters from **Fig. 1I**. The size of the circles corresponds to the percentage of cells within the cluster that express the gene. The color corresponds to the average gene expression in those cells compared to the population as a whole. (**C**) Publicly available bulk RNA-seq data from TCGA were used to investigate *CD97* expression in GBM from all TCGA-defined transcriptional subtypes (classical, mesenchymal, and proneural). The neural GBM subtype is outdated and likely includes mixed samples of the other three subtypes and normal brain tissue. (**D**) Table displaying CD97 splice variant (SV) 1-3 full-length and fragment sizes. (**E**) Immunoblots showing glycosylated and deglycosylated bands for the CD97 ICD (equivalent to the CTF) and the ECD (equivalent to NTF). CD97 S531A is an uncleavable point mutant of splice variant 1. Endogenous CD97 expression in the PDGC is also shown (indicated by the black arrow). Lanes have been rearranged for clarity. Non-specific bands are indicated by a # sign.

**Supplementary Figure 2: CD97 ablation impacts PDGC proliferation and viability.** (**A**) Schematic of the *ADGRE5* gene showing exon targeting of the shRNA and gRNA constructs used in this paper. Corresponding domains are shown beneath the gene. Note gRNA2 targets an EGF domain not found in CD97 splice variant 2. (**B**) Histogram showing CD97 surface expression in PDGCs infected with the gRNA construct (mCherry+) and uninfected PDGCs (mCherry-) 20 days post-infection (dpi). Reduced CD97 surface expression is specific to PDGCs infected with a gRNA targeting the *ADGRE5* gene and is not observed in those cells infected with a control gRNA targeting the human homologue of the *ROSA26* locus. (**C**) Fluorescent images of DAPI-stained PDGCs in the upper (total) and lower (migrated) chamber of a transwell assay following lentiviral infection with a shRNA against CD97 or a SCR control. A bar graph quantifies the ratio of cells within the lower chamber to cells within the upper chamber [n=6 (two PDGCs compiled); paired t-test; *, $p<0.05$]. (**D**) A bar graph quantifying the number of tumorspheres in a tumorsphere formation assay after doxycycline-induced CD97 overexpression of splice variants 1-3. The graph is normalized to non-induced cells (n=3-4 per splice variant; 2-way ANOVA $F_{1,18}=1.125$; ns, $p>0.05$). (**E**) Bar graphs quantifying the percentage of PDGCs or astrocytes within the S/G2 phase of the cell cycle following CD97 knockdown as measured by Hoechst 33342 staining. Graphs are normalized to a SCR shRNA control. Representative histograms are displayed as well. An arrow indicates the peak with cells in the S/G2 phase of the cell cycle. (GBM: n=2-3 per PDGC; 2-way

ANOVA $F_{1,13}$=5.282, $p<0.05$; NHAs: n=3 unpaired t-test; ns, $p>0.05$). (**F**) Immunofluorescent images of Ki67-stained PDGCs following CD97 knockdown. The percentage of Ki67+ cells are quantified in the bar graph, which is normalized to the SCR shRNA control. (n=2-3 per PDGC; 2-way ANOVA $F_{1,12}$=47.47, $p<0.0001$). (**G**) The percentage of PDGCs or astrocytes that are either Annexin-V+ and/or DAPI+ after CD97 knockdown are quantified in the bar graphs, which are normalized to the SCR control. Representative dot plots are shown with quadrants labeling cells considered viable and non-viable. (GBM: n=3-4 per PDGC; 2-way ANOVA $F_{1,22}$=5.829, $p<0.05$; NHAs: n=3; unpaired t-test; ns, $p>0.05$). Error bars throughout this figure indicate SEM.

**Supplementary Figure 3: Additional metabolic data.** (**A**) Immunoblot staining against GLUT3, GAPDH, and CD97 of whole cell lysates collected from PDGCs following lentiviral transduction with a SCR shRNA, CD97 shRNA, or tet-inducible CD97 overexpression construct in the absence or presence of doxycycline. (**B**) GSEA was used to identify pathway enrichment of transcriptomic data from single replicates of three separate PDGCs following CD97 knockdown. Some of the top pathways enriched after knockdown included pathways related to cellular respiration (red). Some of the depleted pathways after CD97 knockdown included glycolytic pathways (blue). Pathways are ranked based on their normalized enrichment score (NES). (**C**) Representative GSEA plots of pathways from panel 3B generated using GSEA software. (**D**) A heatmap quantifying levels of heavy-carbon intermediates of glycolysis and the TCA-cycle after PDGCs were exposed to heavy-carbon glucose. The flux metabolomic assay includes three biological replicates from the same PDGC following CD97 (or SCR) knockdown. The number of heavy-carbons are listed following the metabolite. All glycolytic metabolite data was generated using PDGCs after 30 minutes of exposure. All TCA cycle metabolite data was generated using PDGCs after 120 minutes of exposure and are organized into passes based on the theoretical number of heavy-carbons incorporated from the heavy-carbon glucose. (**E**) A heatmap quantifying steady-state levels of metabolites relevant for PPP following CD97 knockdown.

**Supplementary Figure 4: Additional signaling data.** (**A**) An immunoblot demonstrating increased levels of ERK1/ERK2 phosphorylation following stable transfection of PDGCs with the $MEK^{DD}$ construct. The bar graph quantifies the densitometry ratios and is normalized to the empty vector control (n=3; paired t-test; **, $p<0.01$). (**B**) Summary statistics for luminescent signal after PDGCs were induced for overexpression of CD97 or an empty vector control and transfected with luciferase reporters for G protein-coupled pathways (CRE, $G\alpha_s$; SRE, $G\beta\gamma$; SRF-RE, $G\alpha_{12/13}$; NFAT, $G\alpha_q$) (n=2-8; unpaired t-tests; ns, $p>0.05$). Data is normalized to uninduced controls. (**C**)

Pull-down assay for GTP-bound RHOA after overexpression of WT CD97 or an EV control. A positive (+) and negative (-) control were included as described by the manufacturer. Lanes are rearranged for clarity. (**D**) Immunoblot for AKT and ERK1/ERK2 and their phosphorylated counterparts after overexpression of CD97 or an empty vector control along with the corresponding densitometry ratios displayed in the bar graph. (**E**) A heatmap showing detection of the five novel phosphorylation sites in the CPTAC dataset collected from 99 GBM samples. Values are based on a global-centered normalization. White boxes indicate no detection in that sample. (**F**) Correlation matrix summarizing the correlation between the five novel phosphorylation sites identified from the publicly available CPTAC phosphoproteomic data collected from 99 GBM samples. (**G**) Sanger sequencing result confirming removal of the ∆PS sequence shown in **Fig. 5B**. (**H**) Immunofluorescent staining against the CD97 N-terminus shows proper membrane localization of WT CD97 and the ∆PS mutant in HEK293 cells. No signal is observed upon transduction with an empty vector control. Error bars throughout the figure indicate SEM.

**Supplementary Figure 5: Additional data on putative ligands.** Publicly available single-cell RNA-seq data of adult and pediatric GBM from the Broad Institute Single-Cell Portal (GSE131928) were used.[1] (**A**) Clusters, identifiers, and the number of cells are shown. (**B**) Transcript levels of *ADGRE5* are shown. (**C,D**) Transcript levels of putative CD97 ligands: *CD55* (**C**) and *THY1/CD90* (**D**) are shown. Our own integrated single-cell data from IDH-wildtype GBM and IDH-mutant astrocytoma (**Fig. 1I**) also shows *ADGRE5* (**E**), *CD55* (**F**), and *THY1/CD90* (**G**) expression.

**Supplementary Figure 6: Additional dose-response curve data.** (**A**) The human anti-CD97 antibody was conjugated to a pH sensitive dye (pHAb) and internalization was quantified via flow cytometry. A non-specific IgG isotype control was included (n=3; paired t-test; ***, p<0.001; ****, p<0.0001). (**B-E**) Individual dose-response curves based on WST8 viability assays in (**B**) NHAs, (**C**) proneural GBM, (**D**) NSCs, and (**E**) classical GBM, after treatment with the CD97 ADC, MMAF (impermeant cytotoxic drug) alone, or MMAE (membrane-permeant cytotoxic drug) alone (n=3 biological replicates each based on three technical triplicates). Curves were based on a nonlinear regression fit. (**F**) A table displaying the $LD_{50}$ values (in nM) for the CD97 ADC, MMAF, and MMAE on all four cell lines as calculated from dose-response curves in **Supp. Fig. 6B-E**. Curves were based on a nonlinear regression fit. Error bars throughout this figure indicate SEM.

**SUPPLEMENTARY METHODS**

**Steady-state and flux metabolomics**

*Extraction of metabolites from Plasma* – Prior to extraction, samples were moved from -80°C storage to wet ice and thawed. Extraction buffer, consisting of 80% methanol (Fisher Scientific) and 500 nM metabolomics amino acid mix standard (Cambridge Isotope Laboratories), was prepared and placed on dry ice. Samples were extracted by mixing 50 µL of sample with 950 µL of extraction buffer in 2.0 mL screw cap vials containing ~100 µL of disruption beads (Research Products International). Each sample was homogenized for 10 cycles on a bead blaster homogenizer (Benchmark Scientific). Cycling consisted of a 30 sec homogenization time at 6 m/s followed by a 30 sec pause. Samples were subsequently spun at 21,000 g for 3 minutes at 4°C. A set volume of each (450 µL) was transferred to a 1.5 mL tube and dried down by speedvac (Thermo Fisher). Samples were reconstituted in 50 µL of Optima LC/MS grade water (Fisher Scientific). Samples were sonicated for 2 minutes, then spun at 21,000 g for 3 minutes at 4°C. 20 µL were transferred to LC vials containing glass inserts for analysis. The remaining sample was placed in -80°C for long term storage.

*LC-MS/MS with the hybrid metabolomics method* – Samples were subjected to an LC-MS analysis to detect and quantify known peaks. A metabolite extraction was carried out on each sample based on a previously described method.[2] The LC column was a Millipore™ ZIC-pHILIC (2.1 x150 mm, 5 µm) coupled to a Dionex Ultimate 3000™ system and the column oven temperature was set to 25°C for the gradient elution. A flow rate of 100 µL/min was used with the following buffers; A) 10 mM ammonium carbonate in water, pH 9.0, and B) neat acetonitrile. The gradient profile was as follows; 80-20% B (0-30 minutes), 20-80% B (30-31 minutes), 80-80% B (31-42 minutes). Injection volume was set to 2 µL for all analyses (42 minutes total run time per injection). MS analyses were carried out by coupling the LC system to a Thermo Q Exactive HF™ mass spectrometer operating in heated electrospray ionization mode (HESI). Method duration was 30 minutes with a polarity switching data-dependent Top 5 method for both positive and negative modes. Spray voltage for both positive and negative modes was 3.5 kV and capillary temperature was set to 320°C with a sheath gas rate of 35, aux gas of 10, and max spray current of 100 µA. The full MS scan for both polarities utilized 120,000 resolution with an AGC target of 3e6 and a maximum IT of 100 ms, and the scan range was from 67-1000 m/z. Tandem MS spectra for both positive and negative mode used a resolution of 15,000, AGC target of 1e5, maximum IT of 50 ms, isolation window of 0.4 m/z, isolation offset of 0.1 m/z, fixed first mass of 50 m/z, and

3-way multiplexed normalized collision energies (nCE) of 10, 35, 80. The minimum AGC target was 1e4 with an intensity threshold of 2e5. All data were acquired in profile mode.

*Relative quantification of metabolites* – The resulting ThermoTM RAW files were converted to mzXML format using ReAdW.exe version 4.3.1 to enable peak detection and quantification. The centroided data were searched using an in-house python script Mighty_skeleton version 0.0.2 and peak heights were extracted from the mzXML files based on a previously established library of metabolite retention times and accurate masses adapted from the Whitehead Institute[3], and verified with authentic standards and/or high-resolution MS/MS spectral manually curated against the NIST14MS/MS[4] and METLIN (2017)[5] tandem mass spectral libraries. Metabolite peaks were extracted based on the theoretical m/z of the expected ion type e.g., [M+H]+, with a ±5 part-per-million (ppm) tolerance, and a ± 7.5 second peak apex retention time tolerance within an initial retention time search window of ± 0.5 minutes across the study samples. The resulting data matrix of metabolite intensities for all samples and blank controls was processed with an in-house statistical pipeline Metabolyze version 1.0 and final peak detection was calculated based on a signal to noise ratio (S/N) of 3X compared to blank controls, with a floor of 10,000 (arbitrary units). For samples where the peak intensity was lower than the blank threshold, metabolites were annotated as not detected, and the threshold value was imputed for any statistical comparisons to enable an estimate of the fold change as applicable. The resulting blank corrected data matrix was then used for all group-wise comparisons, and t-tests were performed with the Python SciPy (1.1.0)[6] library to test for differences and generate statistics for downstream analyses. Any metabolite with p-value < 0.05 was considered significantly regulated (up or down).

1 Neftel, C. *et al.* An Integrative Model of Cellular States, Plasticity, and Genetics for Glioblastoma. *Cell* **178**, 835-849.e821, doi:10.1016/j.cell.2019.06.024 (2019).
2 Jones, D. R., Wu, Z., Chauhan, D., Anderson, K. C. & Peng, J. A nano ultra-performance liquid chromatography-high resolution mass spectrometry approach for global metabolomic profiling and case study on drug-resistant multiple myeloma. *Anal Chem* **86**, 3667-3675, doi:10.1021/ac500476a (2014).
3 Chen, W. W., Freinkman, E., Wang, T., Birsoy, K. & Sabatini, D. M. Absolute Quantification of Matrix Metabolites Reveals the Dynamics of Mitochondrial Metabolism. *Cell* **166**, 1324-1337.e1311, doi:10.1016/j.cell.2016.07.040 (2016).
4 Simón-Manso, Y. *et al.* Metabolite profiling of a NIST Standard Reference Material for human plasma (SRM 1950): GC-MS, LC-MS, NMR, and clinical laboratory analyses, libraries, and web-based resources. *Anal Chem* **85**, 11725-11731, doi:10.1021/ac402503m (2013).
5 Smith, C. A. *et al.* METLIN: a metabolite mass spectral database. *Ther Drug Monit* **27**, 747-751, doi:10.1097/01.ftd.0000179845.53213.39 (2005).

6 Jones, E., Oliphant, T. & Peterson, P. SciPy: Open Source Scientific Tools for Python. (2001).

A

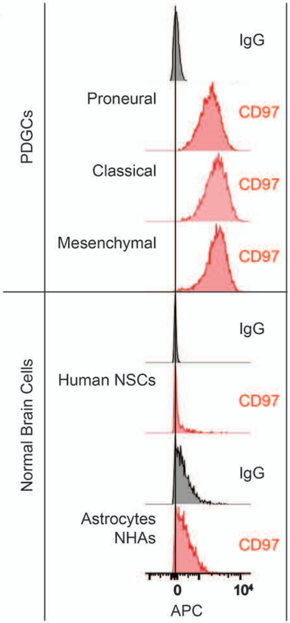


B

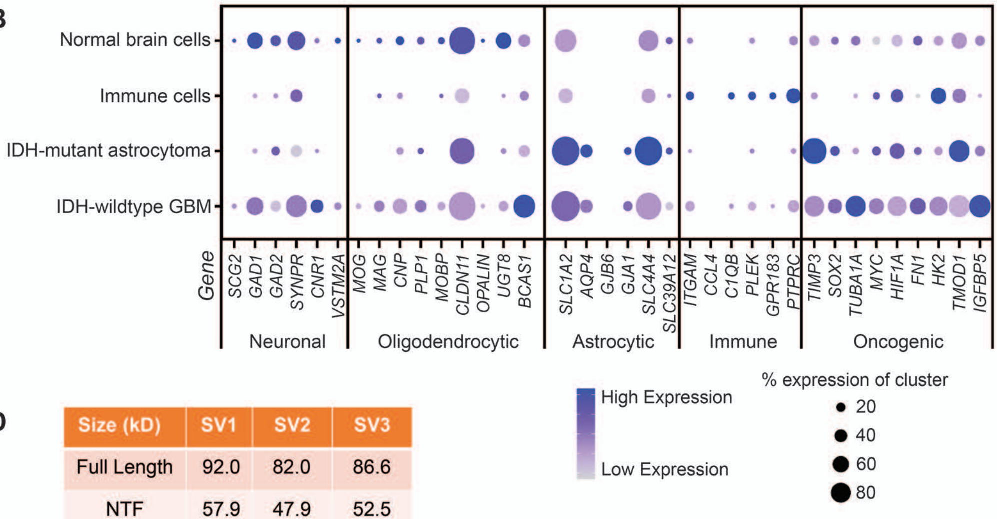


D

| Size (kD) | SV1 | SV2 | SV3 |
|---|---|---|---|
| Full Length | 92.0 | 82.0 | 86.6 |
| NTF | 57.9 | 47.9 | 52.5 |
| CTF | 34.1 | 34.1 | 34.1 |

C

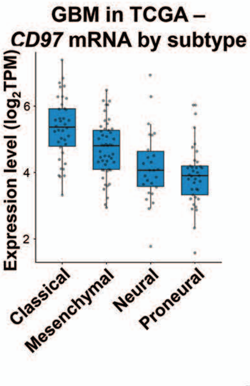


E

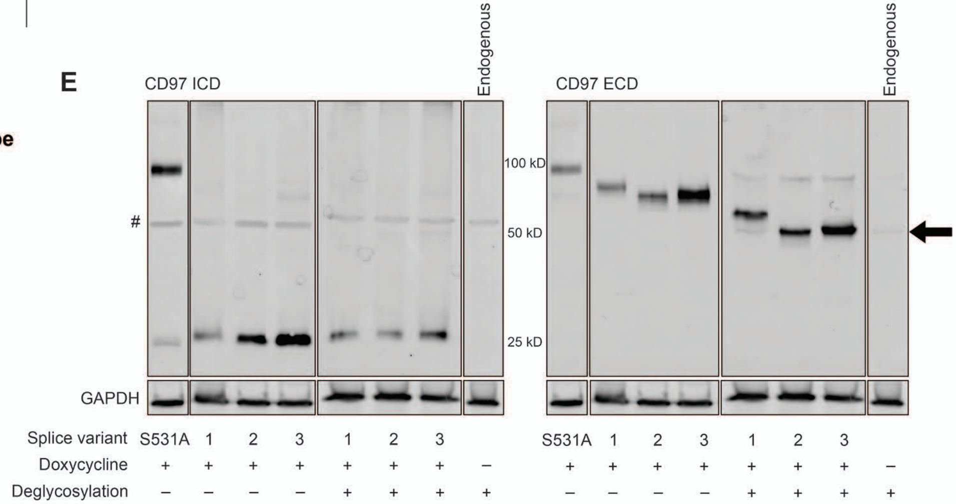

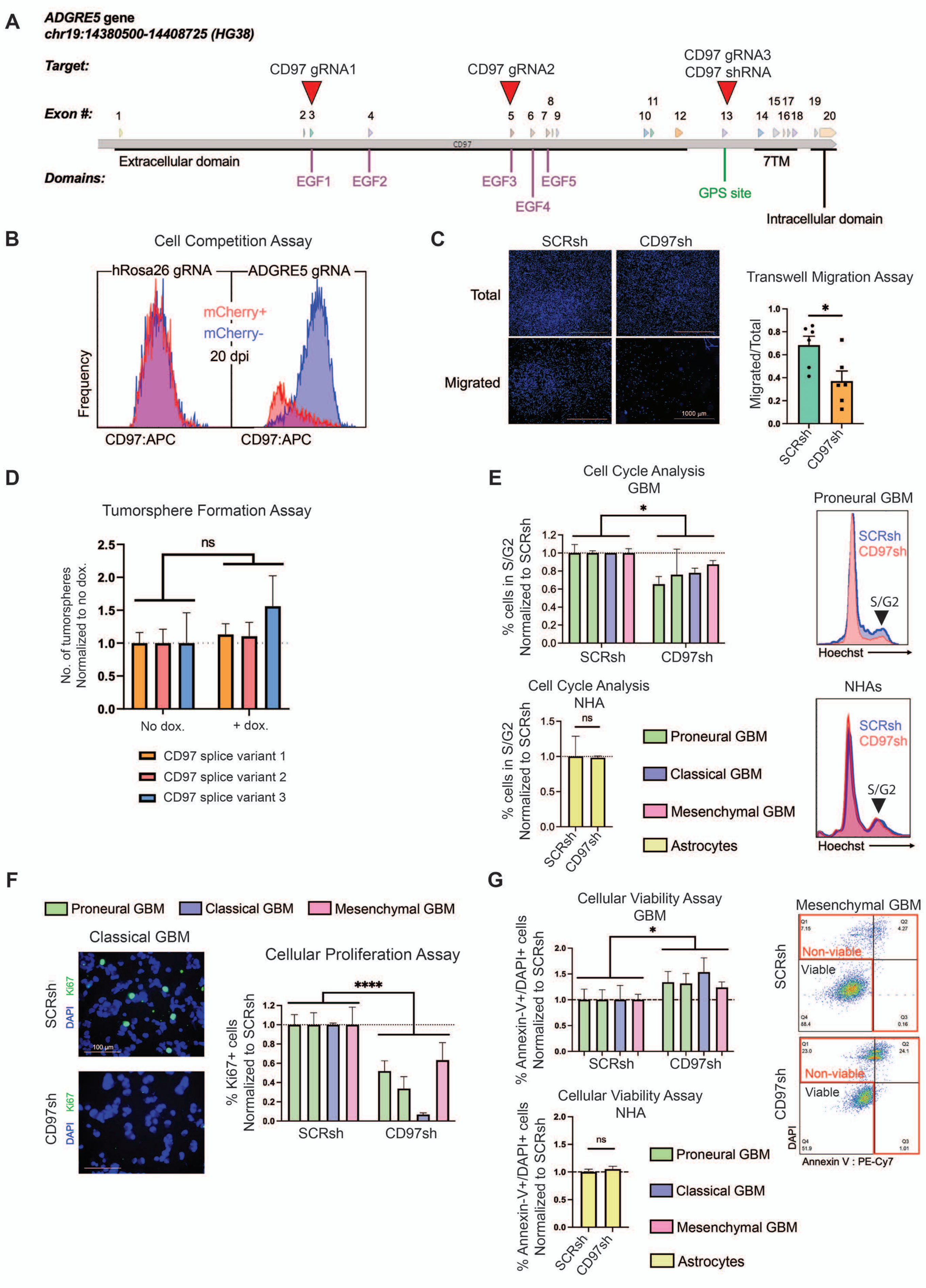
A
ADGRE5 gene
chr19:14380500-14408725 (HG38)
Target:
CD97 gRNA1
CD97 gRNA2
CD97 gRNA3
CD97 shRNA
Exon #:
Extracellular domain
7TM
Domains:
EGF1
EGF2
EGF3
EGF4
EGF5
GPS site
Intracellular domain
B
Cell Competition Assay
hRosa26 gRNA
ADGRE5 gRNA
mCherry+
mCherry-
20 dpi
Frequency
CD97:APC
CD97:APC
C
SCRsh
CD97sh
Total
Migrated
1000 µm
Transwell Migration Assay
Migrated/Total
SCRsh
CD97sh
D
Tumorsphere Formation Assay
ns
No. of tumorspheres Normalized to no dox.
No dox.
+ dox.
CD97 splice variant 1
CD97 splice variant 2
CD97 splice variant 3
E
Cell Cycle Analysis GBM
% cells in S/G2 Normalized to SCRsh
SCRsh
CD97sh
Proneural GBM
SCRsh
CD97sh
S/G2
Hoechst
Cell Cycle Analysis NHA
ns
NHAs
Proneural GBM
Classical GBM
Mesenchymal GBM
Astrocytes
F
Proneural GBM
Classical GBM
Mesenchymal GBM
Classical GBM
SCRsh
CD97sh
Ki67
DAPI
100 µm
Cellular Proliferation Assay
****
% Ki67+ cells Normalized to SCRsh
G
Cellular Viability Assay GBM
% Annexin-V+/DAPI+ cells Normalized to SCRsh
Cellular Viability Assay NHA
Mesenchymal GBM
Non-viable
Viable
DAPI
Annexin V : PE-Cy7

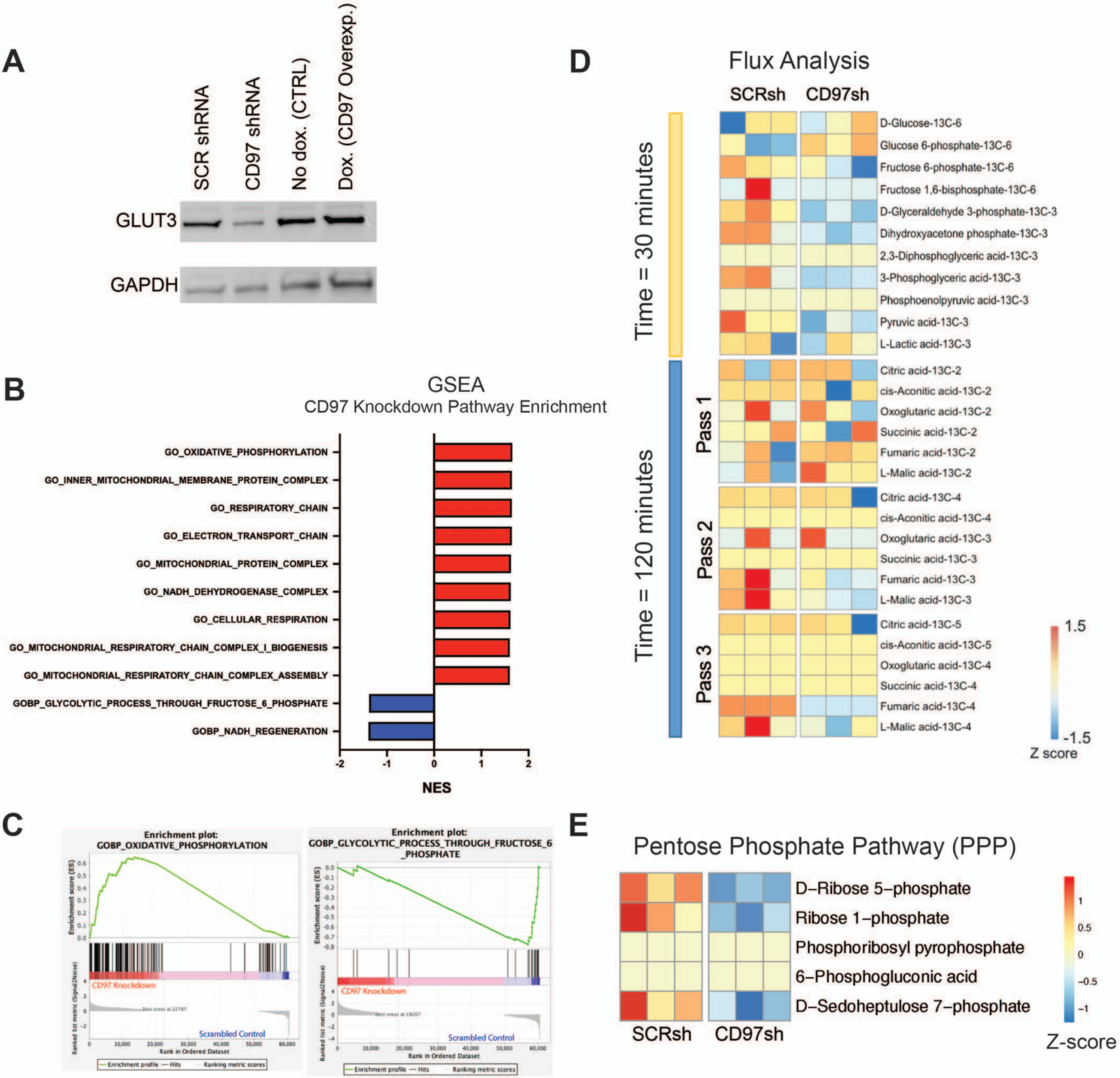

A
SCR shRNA
CD97 shRNA
No dox. (CTRL)
Dox. (CD97 Overexp.)
GLUT3
GAPDH
B
GSEA
CD97 Knockdown Pathway Enrichment
GO_OXIDATIVE_PHOSPHORYLATION
GO_INNER_MITOCHONDRIAL_MEMBRANE_PROTEIN_COMPLEX
GO_RESPIRATORY_CHAIN
GO_ELECTRON_TRANSPORT_CHAIN
GO_MITOCHONDRIAL_PROTEIN_COMPLEX
GO_NADH_DEHYDROGENASE_COMPLEX
GO_CELLULAR_RESPIRATION
GO_MITOCHONDRIAL_RESPIRATORY_CHAIN_COMPLEX_I_BIOGENESIS
GO_MITOCHONDRIAL_RESPIRATORY_CHAIN_COMPLEX_ASSEMBLY
GOBP_GLYCOLYTIC_PROCESS_THROUGH_FRUCTOSE_6_PHOSPHATE
GOBP_NADH_REGENERATION
-2
-1
0
1
2
NES
C
Enrichment plot:
GOBP_OXIDATIVE_PHOSPHORYLATION
Enrichment score (ES)
CD97 Knockdown
Scrambled Control
Ranked list metric (Signal2Noise)
Rank in Ordered Dataset
Enrichment profile
Hits
Ranking metric scores
Enrichment plot:
GOBP_GLYCOLYTIC_PROCESS_THROUGH_FRUCTOSE_6_PHOSPHATE
Enrichment score (ES)
CD97 Knockdown
Scrambled Control
Ranked list metric (Signal2Noise)
Rank in Ordered Dataset
Enrichment profile
Hits
Ranking metric scores
D
Flux Analysis
SCRsh
CD97sh
Time = 30 minutes
Time = 120 minutes
Pass 1
Pass 2
Pass 3
D-Glucose-13C-6
Glucose 6-phosphate-13C-6
Fructose 6-phosphate-13C-6
Fructose 1,6-bisphosphate-13C-6
D-Glyceraldehyde 3-phosphate-13C-3
Dihydroxyacetone phosphate-13C-3
2,3-Diphosphoglyceric acid-13C-3
3-Phosphoglyceric acid-13C-3
Phosphoenolpyruvic acid-13C-3
Pyruvic acid-13C-3
L-Lactic acid-13C-3
Citric acid-13C-2
cis-Aconitic acid-13C-2
Oxoglutaric acid-13C-2
Succinic acid-13C-2
Fumaric acid-13C-2
L-Malic acid-13C-2
Citric acid-13C-4
cis-Aconitic acid-13C-4
Oxoglutaric acid-13C-3
Succinic acid-13C-3
Fumaric acid-13C-3
L-Malic acid-13C-3
Citric acid-13C-5
cis-Aconitic acid-13C-5
Oxoglutaric acid-13C-4
Succinic acid-13C-4
Fumaric acid-13C-4
L-Malic acid-13C-4
1.5
-1.5
Z score
E
Pentose Phosphate Pathway (PPP)
D−Ribose 5−phosphate
Ribose 1−phosphate
Phosphoribosyl pyrophosphate
6−Phosphogluconic acid
D−Sedoheptulose 7−phosphate
SCRsh
CD97sh
1
0.5
0
−0.5
−1
Z-score

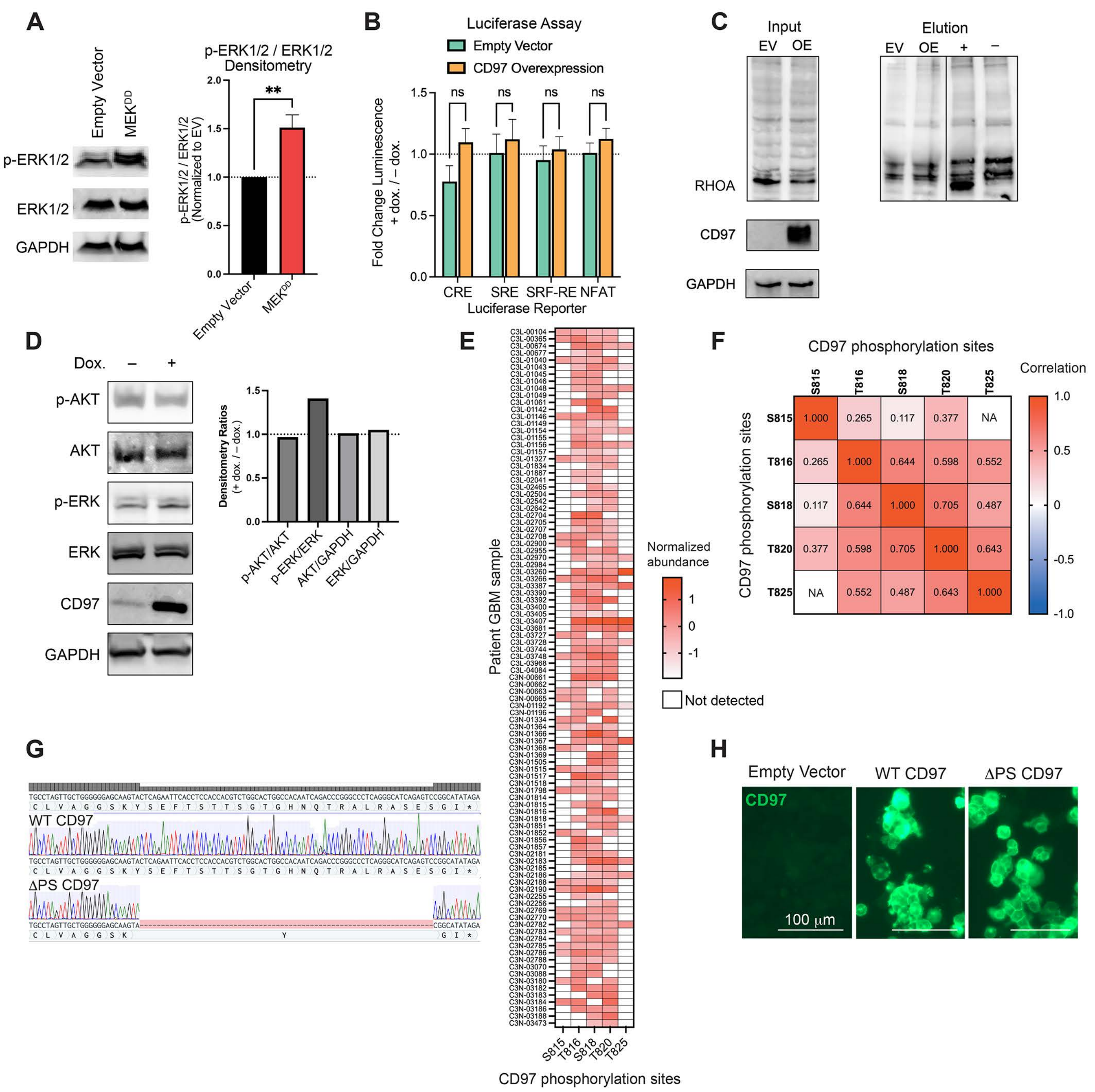
A
Empty Vector
MEK^DD
p-ERK1/2
ERK1/2
GAPDH
p-ERK1/2 / ERK1/2 Densitometry
p-ERK1/2 / ERK1/2 (Normalized to EV)
**
B
Luciferase Assay
Empty Vector
CD97 Overexpression
Fold Change Luminescence + dox. / – dox.
ns
CRE
SRE
SRF-RE
NFAT
Luciferase Reporter
C
Input
Elution
EV
OE
+
–
RHOA
CD97
GAPDH
D
Dox.
p-AKT
AKT
p-ERK
ERK
CD97
GAPDH
Densitometry Ratios (+ dox. / – dox.)
p-AKT/AKT
p-ERK/ERK
AKT/GAPDH
ERK/GAPDH
E
Patient GBM sample
Normalized abundance
Not detected
S815
T816
S818
T820
T825
CD97 phosphorylation sites
F
CD97 phosphorylation sites
Correlation
G
WT CD97
ΔPS CD97
H
Empty Vector
WT CD97
ΔPS CD97
CD97
100 μm

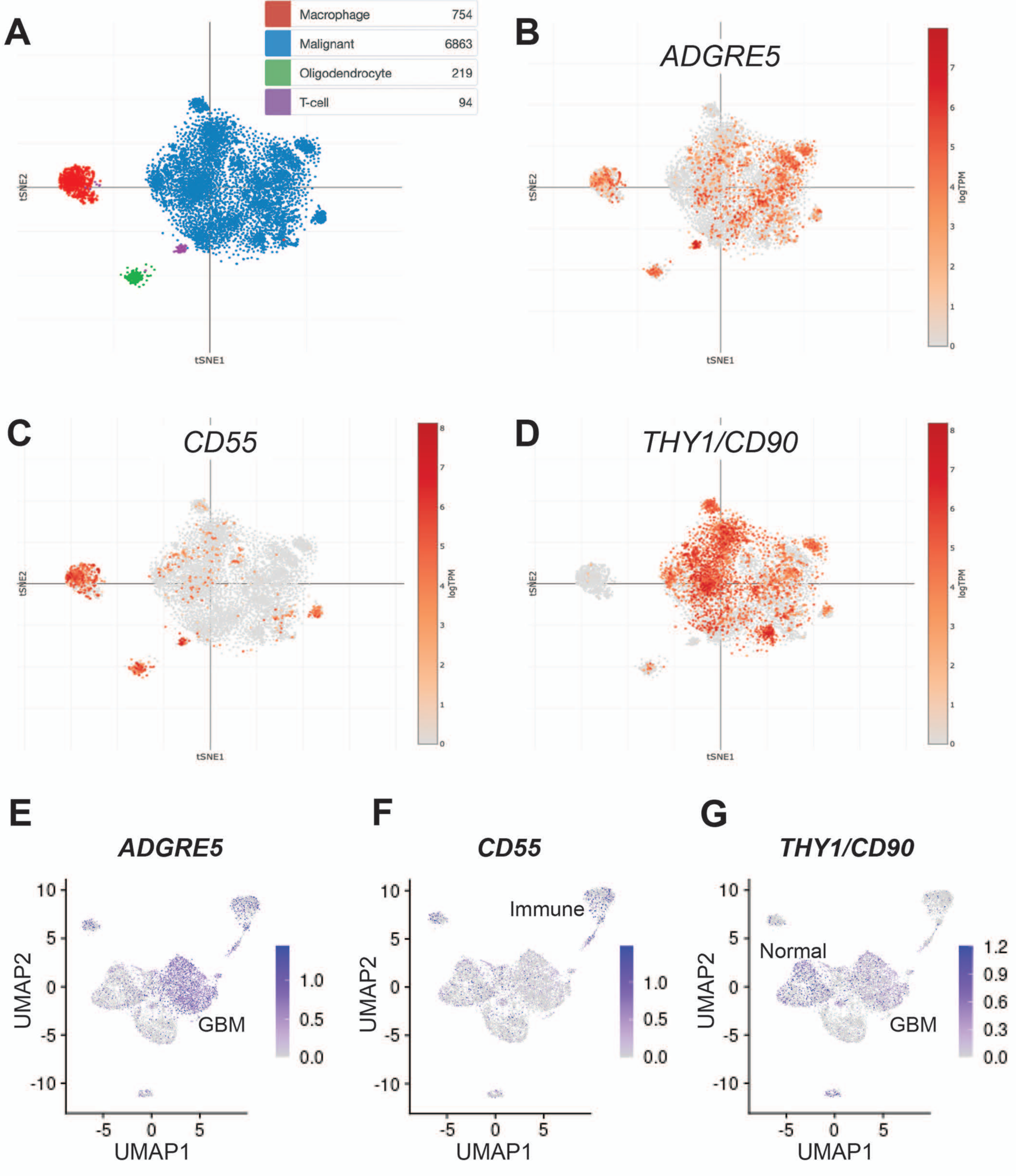
A
Macrophage 754
Malignant 6863
Oligodendrocyte 219
T-cell 94
tSNE1
tSNE2
B
ADGRE5
logTPM
C
CD55
D
THY1/CD90
E
ADGRE5
UMAP1
UMAP2
GBM
F
CD55
Immune
G
THY1/CD90
Normal
GBM

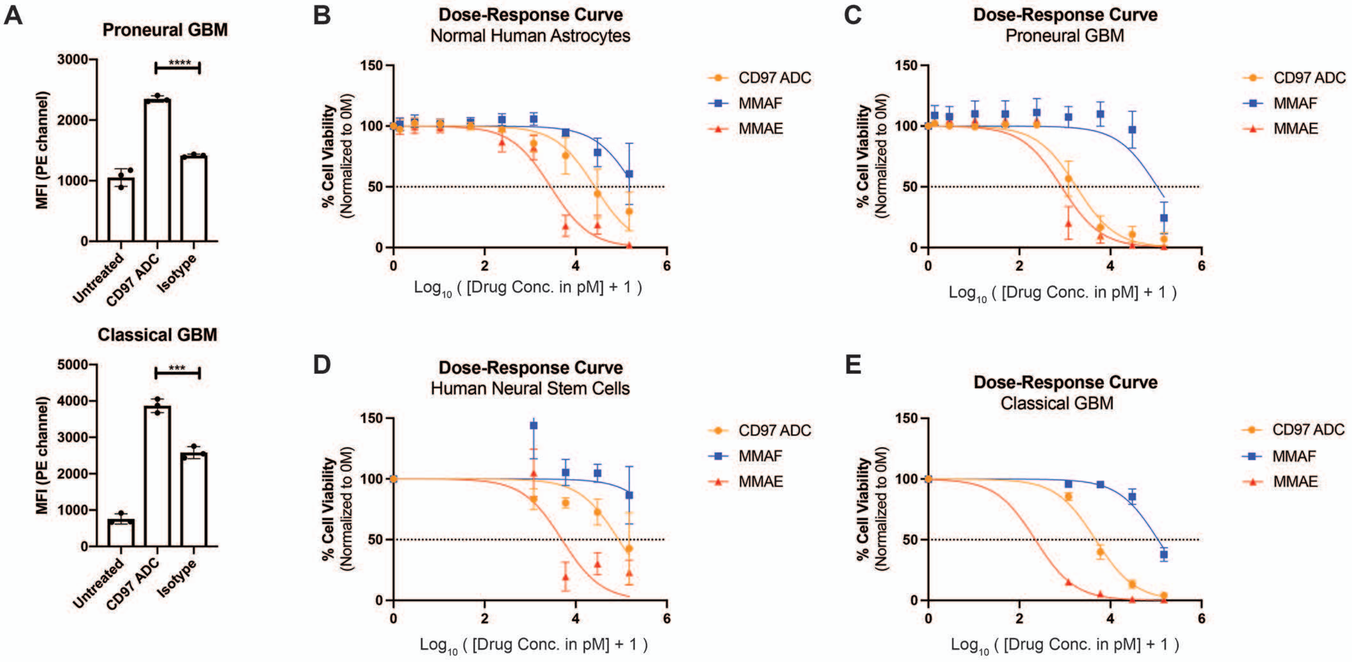


F

$LD_{50}$ (nM)

| Cell Line | CD97 ADC |
|---|---|
| NHAs | 26.27 |
| Proneural | 1.78 |
| NSCs | 87.30 |
| Classical | 4.70 |

$LD_{50}$ (nM)

| Cell Line | MMAF | MMAE |
|---|---|---|
| NHAs | 196.37 | 2.81 |
| Proneural | 106.83 | 0.84 |
| NSCs | 1212.30 | 4.98 |
| Classical | 110.55 | 0.22 |
